# Experimental conditions to retrieve intrinsic cooperativity α directly from single binding assay data exemplified by the ternary complex formation of FKBP12, MAPRE1 and macrocyclic molecular glues

**DOI:** 10.1101/2025.01.15.631501

**Authors:** Jan Schnatwinkel, Richard R. Stein, Michael Salcius, Julian Wong, Shu-Yu Chen, Marianne Fouché, Hans-Joerg Roth

## Abstract

The incorporation of disease-relevant targets in a compound-dependent manner into ternary complexes with an assisting chaperone protein has become a common modality. Among the various types of ternary complex-forming compounds, molecular glues are of particular interest because they do not require significant binary (independent) target affinity. Instead, ternary complexes formed by molecular glues retrieve a significant part of their thermodynamic stability through newly induced chaperone–target or glue–target interactions that occur only in the ternary complex. These interactions lead to enhanced ligand binding – termed intrinsic cooperativity α – which can be retrieved via the apparent cooperativity either by monitoring ligand binding through the chaperone or through the target protein. In this publication, the advantage of measuring the apparent cooperativity to determine the cooperativity α by the weaker binding protein is discussed and illustrated using the example of ternary complexes between FKBP12, MAPRE1 and macrocyclic molecular glues derived from the rapamycin binding motif for FKBP12. Furthermore, the impact of the following three parameters on the apparent cooperativity is illustrated: 1.) the concentration of the monitoring protein, 2.) the excess of the counter protein, 3.) the affinity of the glue to the weaker binding protein in combination with the degree of intrinsic cooperativity α. From this, experimental conditions to determine the intrinsic cooperativity α with only one binding assay and without the need for a comprehensive mathematical model covering all simultaneous events under non-saturating conditions are highlighted. However, this framework requires a binding assay capable of measuring or at least estimating very weak binary affinities. If this is not possible for experimental reasons, but binding assays for both proteins are available within a normal bandwidth and the affinity to the stronger binding protein is not too high, it is discussed how the binding curve for the weaker binding protein in the presence of an excess of the weaker binding protein can be used to overcome the missing binary *K*_d_ for the weakly binding protein.

## Introduction

Bringing disease-relevant targets in a compound-dependent manner into proximity to a second (assisting) protein to enable the formation of a ternary complex has become a common modality^1-8^. In addition to simply inhibiting the function of target proteins by forming such ternary complexes, many other induced proximity modalities have emerged in the recent past. Depending on the nature of the second protein, often also referred to as “chaperone” protein, target proteins can undergo an induced change in location or undergo chemical modifications such as ubiquitination (to trigger degradation), de-ubiquitination (to rescue from degradation), phosphorylation, de-phosphorylation (to modulate signalling cascades) or other enzymatic transformations.^9-14^

While bifunctional ternary complex-forming compounds require an independent binary affinity to both, the target and the chaperone protein, this is not the case for molecular glues. They do not have necessarily significant affinity to either protein or, more often, to only one of the two proteins.^15-17^ This makes molecular glues attractive as modulating agents for targets where it is difficult to develop chemical matter with a high independent binary binding affinity. While a small molecule compound alone is not able to bind tightly enough to flat, large and floppy surfaces to modulate the function of such targets, the surface of a pre-formed binary complex consisting of the molecular glue and the chaperone protein – depending on the degree of cooperativity – can be able to do so.^16, 18^ This mode of action also offers a high potential for selectivity, even isoform selectivity, as a much larger surface area of the target is involved in a ternary complex and subtle changes remote from often conservative low molecular weight binding sites still can have a chance to contribute to differences in the binding affinity of the pre-formed molecular glue–chaperone complex to the target.^19-28^ Molecular glues also tend – with increasing cooperativity α – to have a lower molecular weight than bifunctional compounds with the same thermodynamic stability of the ternary complex. These two properties, i.e., low molecular weight and high selectivity make molecular glues to attractive hit and lead candidates.^29^

In our recent publication, we highlighted the important difference between apparent cooperativity, an EC_50_ shift that is highly dependent on experimental parameters, and the intrinsic cooperativity α, which is a natural constant of a compound for a given ternary complex that is independent of any experimental parameters. Like a *K*_d_, the intrinsic cooperativity α is therefore ideally suited for compound characterization. We have also shown through simulations, i.e., from a theoretical point of view, that it is advantageous to characterize cooperativity by monitoring binding without and with presence of the counter protein through the weaker binding protein, which is often the target. The reason for this is that apparent cooperativity values, i.e., the EC_50_ shifts of a measured molecular glue binding curve in the absence and presence of the counter protein – required experimental values to retrieve the intrinsic cooperativity α – are much higher than when measured through the stronger binding protein. The smaller these shifts are, the lower the signal-to-noise ratio is. If, for example, a molecular glue has a binary affinity to the chaperone of less than 100 nM and the monitoring method requires chaperone concentrations of 20 nM, for reasons of sensitivity, the assay will likely bottom-out and is, hence, not suitable for estimating the intrinsic cooperativity α via shifts in the apparent cooperativity or EC_50_ values in an acceptable signal-to-noise ratio.^29^

In this work, we present binding and cooperativity studies of a small series of molecular glues which form ternary complexes consisting of FKBP12, the corresponding macrocyclic glues and the plus-end tracking protein (+TIPs) MAPRE1. We experimentally demonstrate the advantage of monitoring cooperativity through the weaker binding protein MAPRE1, emphasizing the value of methods that can accurately measure very low binding affinities. Furthermore, we discuss the experimental conditions, which are required to coincide the EC_50_ value of the %T_bound, ternary_ curve (i.e., the normalized ligand binding to the target, %T_bound, ternary_=([TL]+[TLC])/[T_tot_]·100%, in presence of an excess of the assisting protein)^29^ with the *K*_T,2_ of the corresponding ternary complex, which leads finally to conditions where the intrinsic cooperativity α can be retrieved only from the assay for the weaker binding protein instead of from two binding assays.

### Retrieving cooperativity α from binding measurements to one instead of two proteins

#### The difference between apparent cooperativity and intrinsic cooperativity α

The relationship between the intrinsic cooperativity α and the apparent cooperativity can be compared to the relationship between mass and weight, which is why we could call weight also “apparent mass”. Mass is an absolute and intrinsic property of objects and is always the same regardless of the environment although – like all absolute values – it is difficult to measure. Much easier to measure is weight – a relative value – which represents the effect of an object’s mass under certain experimental conditions. Similarly, the apparent cooperativity is the effect of an underlying absolute, intrinsic cooperativity α concealed by specific experimental conditions, i.e., the applied protein concentrations, more specifically, the ratio of target and chaperone protein concentrations relative to the corresponding *K*_C,1_ and *K*_T,1_ values (Fig. 4). Since the apparent cooperativity is the quotient of two EC_50_ values and the intrinsic cooperativity α is the quotient of two *K*_d_ values (equation 1), the difference between apparent and intrinsic cooperativity α is the result of the difference in EC_50_ and *K*_d_ values. The EC_50_ value describes the resulting effect of the underlying absolute and intrinsic affinity of a ligand to a protein, i.e., the *K*_d_ value adjusted by the applied protein concentration, refers to the applied ligand concentration ([L_tot_]) required to achieve 50% of this effect. For this reason, EC_50_ values could also be referred to as “apparent *K*_d_”. In contrast, the *K*_d_ is an absolute intrinsic property of a protein–ligand pair and, hence, independent of the experimental conditions and always refers to concentrations in free ligand, [L].

#### The general requirement of having measured *K*_d_s referring to both proteins to determine the cooperativity α

In our previous publication, we presented a simulation tool that enabled the determination of the cooperativity α by iteratively comparing the experimentally measured EC_50_ value. Specifically, it was determined by matching given [L_tot_] of the %T_bound, ternary_ curve and a numerically estimated [L_tot_] constrained on [L]=[L_tot_]–[CL]–[TL]– [CLT]. To calculate the concentrations of CL, TL and CLT on the basis of [L], the corresponding algebraic expressions are used.^29^ The above-mentioned general model covers ternary complex-forming compounds ranging from bifunctional compounds to molecular glues type I, which share at least one weak binary independent binding affinity for both proteins. The model assumes that these measured *K*_d_ values for *K*_C,1_ and *K*_T,1_ are available to calculate concentration values for CL, TL and CLT as a function of L_tot_ at equilibrium. The case of an infinite weak binding of the molecular glue to one of the two binding partners (molecular glue type II) and the case of an infinite weak binding to both proteins but an intrinsic and significant affinity of the two proteins to each other (molecular glue type III) requires additional mathematical constraints and will be discussed elsewhere.

#### The advantage of measuring apparent cooperativity through the weaker binding protein

Molecular glues have very different binary affinities for the chaperone (*K*_C,1_) and the target protein (*K*_T,1_). A typical pattern is a high affinity for the chaperone (C) and a very weak or even unmeasurable affinity for the target protein (T), which makes the measurement of the intrinsic cooperativity α a challenge and pushes the above-mentioned model to its limits. When it is possible to measure the *K*_d_ referring to the weak affinity, the different affinities for the chaperone and the target protein result in very different apparent cooperativity values depending on whether the EC_50_ shifts are measured through the weaker or the stronger binding protein, although the underlying intrinsic cooperativity α is the same for the experiments in both directions. The weaker the affinity of the molecular glue L to the monitoring protein (e.g., target T) and the higher the underlying cooperativity α, the greater the EC_50_ shift of the %T_bound,ternary_=([TL]+[CLT])/[T_tot_]·100% curve in the presence of the stronger binding counter protein C compared to the same binding experiments in the absence of the counter protein (%T_bound, binary_ =[TL]/[T_tot_]·100%. Conversely, the stronger the intrinsic binary affinity of the molecular glue L to the monitoring chaperone protein C, the smaller the EC_50_ shifts due to the presence of the weaker binding counter protein, even if the underlying intrinsic cooperativity α is very high^29^. Consequently, the measurement of cooperativity by the stronger binding protein C becomes slightly difficult as the binding assay often cannot distinguish between the strong intrinsic binary ligand binding and the even stronger ternary binding caused by the presence of the counter protein T. In other words, the assay is bottoming out and would require lowering the concentration of the monitored protein C, which is often not possible for sensitivity reasons. Monitoring binary versus ternary binding by the weaker binding protein therefore results in a much better signal-to-noise ratio for the EC_50_ shift, leading directly to more accurate and significant values for apparent cooperativity and ultimately the cooperativity α. But, as mentioned above, the challenge is often to accurately measure the weak affinity.

If the development of an assay for the weaker binding protein T is feasible, for instance, for *K*_T,1_ as high as 100,000 nM, the determination of the missing very weak *K*_T,2_ is possible from the %T_bound, ternary_ curve, which represents under the excess of protein C (e.g., [C_tot_]=10,000 nM) *K*_T,2_. If the ternary complex formation has sufficient intrinsic cooperativity α, the EC_50_ of the corresponding curve should then shift into the assay window for EC_50_ values of less than 100,000 nM and be easily measurable. In such case, the developed mathematical model that requires input values for C_tot_, T_tot_, *K*_C,1_, *K*_T,1_ and α to output ternary complex concentrations [CLT]=*f*(C_tot_, T_tot_, *K*_C,1_, *K*,_T,1_, α) for all cooperativity values α has to be adjusted. According to Equation 1, the missing *K*_T,1_ has to be replaced by *K*_T,2_·α to iteratively match the EC_50_ of the %C_bound,ternary_ curve:

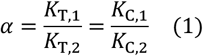

In this case, however, the shift corresponding to the stronger binding protein C derived from the %C_bound, binary_ to the %C_bound, ternary_ curve is required.

#### Retrieving intrinsic cooperativity α by monitoring binding only through one protein

Another advantage of assessing cooperativity through the weaker binding protein is that under specific conditions the EC_50_ of the ternary binding experiment corresponds directly to the *K*_d_ of the ternary complex formation following CL formation (*K*_T,2_). In such cases, assuming that the corresponding binary *K*_T,1_ (which would be necessary to inform the mathematical model requiring both *K*_d_ values) is available, the obtained apparent cooperativity naturally coincides with the intrinsic cooperativity α. Moreover, if such conditions can be identified and are experimentally feasible, the binding assay to the stronger binding protein C is not required and the intrinsic cooperativity α can be determined experimentally by monitoring the binding of a single binding assay only, which relaxes the previously stated requirement of measuring two binary *K*_d_s (*K*_C,1_, *K*_T,2_).^29^

The law of pathway independence states that both ternary complex-forming pathways – the one via CL and the one via TL – result in the same free energy difference between the ternary complex CLT and the sum of the three free species. Consequently, the cooperativity α is the same in both pathways and can therefore be derived from each pathway alone (Equation 1). In other words, the degree to which the presence of T facilitates the ligand binding to C is the same as for the binding of ligand to T in the presence of C relative to the corresponding binary binding. The practical consequence is that the intrinsic cooperativity α can be measured by monitoring the binary and ternary ligand binding to only one of the proteins (T or C) without the use of a mathematical model. The counter protein must be present and controlled with respect to concentration, but no additional binding assay with the counter protein is required.

The problem of retrieving the cooperativity α from a single binding assay is grounded on the question under which conditions the EC_50_s of the %T_bound, binary_ and %T_bound, ternary_ curves correspond to *K*_T,1_ and *K*_T,2_, respectively. More generally, this question can be seen in the context of the difference between EC_50_ and *K*_d_ regardless of whether it is a binary or a ternary experiment.

In a binary binding experiment, *K*_T,1_ corresponds to the concentration of free ligand L ([L]) at the binary inflection point [T] = [TL]:

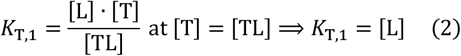

Similarly, at the ternary inflection point, i.e., when [T] = [CLT], *K*_T,2_ corresponds to the concentration of the pre-formed chaperon– ligand complex CL ([CL]):

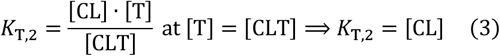

In contrast to the *K*_d_s, the EC_50_ values corresponding to the inflection points of the binary and ternary %T_bound_ curves never coincide with the concentrations of free ligand L ([L]) nor of [CL] but match the concentrations of the total ligand [L_tot_] applied. Specifically, for the binary binding, this means that the EC_50_ value directly corresponds to [L] only if [L_tot_] is equal to [L]. Alternatively, a mathematical relationship between [L_tot_] and [L], i.e., the EC_50_ value of the %T_bound,binary_ curve and the universal *K*_T,1_, which is independent of the applied total concentration of the target protein [T_tot_], would need to be established such that *K*_T,1_=*f*(EC_50_, L_tot_, T_tot_). The same applies to the ternary binding event. Either there are conditions under which the EC_50_ value of the %T_bound, ternary_ match *K*_T,2_ so it can be directly read off the assay data, or a mathematical relationship must be established such that *K*_T,2_ can be derived from the EC_50_ value of the %T_bound, ternary_ curve, i.e., *K*_T,2_=*f*(EC_50_, L_tot_, T_tot_). The minimal benefit is that this mathematical relationship guides the selection of conditions under which the EC_50_ value matches *K*_d_ so that the cooperativity α can be derived directly from the apparent cooperativities from single binding assay data.

In our previous publication^29^, we demonstrated by simulated binding curves that apparent cooperativity values measured through the weaker binding protein are closer to the intrinsic cooperativity α than those measured by the stronger binding protein. Here we specify the exact conditions under which apparent cooperativity measured by the weaker binding protein matches the cooperativity α. Even if only one – namely with the weaker binding protein – and not two binding assays are required, the counter protein as such must be available for the ternary binding experiment, but not as a binary binding assay. This can be a significant advantage in a drug discovery program. One potential obstacle to develop a binary binding assay for the stronger binding protein is the lack of a suitable position for the installation of the tag that reports the binding. Ideally, it should be close to the binding site, but not too close, which is not always possible. An alternative to monitoring binding indirectly is a competitive binding assay with a tag attached to a tracer ligand. However, if the affinity of the tracer ligand is too high, the dynamic range of the assay suffers, which often leads low signal-to-noise ratios. Finally, it is simply less time-consuming and more efficient to determine the intrinsic cooperativity α from only one binding assay than from two, even more so, if the final result is the same.

In the following sections, we discuss three experimental parameters that determine the range in which the EC_50_ value of a ternary binding experiment (from the %T_bound, ternary_ curve) coincides with the ternary *K*_d_ *K*_T,2_. If, additionally, *K*_T,1_ can be measured, the observed apparent cooperativity then coincides with the intrinsic cooperativity α. The three decisive parameters are 1.) the influence of the excess concentration of the stronger binding counter protein, 2.) the role of the concentration of the monitored weaker binding protein, and 3.) the influence of lower *K*_T,1_ on saturation conditions.

#### Conditions under which apparent and intrinsic cooperativity α coincides

##### The influence of the concentration of the stronger binding protein

Measuring ligand binding by monitoring only the changes in the protein to which the ligand binds, so-called target engagement assays, allows a direct comparison between binary and ternary binding experiments through the presence of the counter protein, which is usually unlabelled. The presence of the counter protein causes – in cooperative systems, i.e., with α>1 – additional interactions that only occur in the ternary complex leading to enhanced ligand binding. The assay quantifies this by shifts of the EC_50_ value in the binding curve in the absence (%T_bound, binary_) and in the presence of the counter protein (%T_bound, ternary_). The underlying reason for this EC_50_ shift in the %T_bound, ternary_ curve is a lower *K*_d_ of the ligand in the presence of the counter protein. The intrinsic cooperativity α is defined as the ratio of the binary and the ternary *K*_d_, *K*_T,1_ and *K*_T,2_, respectively. For intrinsic cooperativities greater than 1, the ligand is bound more tightly in the presence of the counter protein. In the absence of cooperativity (α=1), the EC_50_ values of the binary and ternary binding curves are identical and the ligand is not bound more tightly to the target in the presence of the chaperone (Equation 1). This holds regardless of whether ternary complex formation takes place or not. Moreover, this means that a target engagement assay can only detect ternary complex-forming compounds that exhibit intrinsic cooperativity, i.e., α>1 or α<1. For non-cooperative compounds (α=1), an assay that detects the compound-dependent induced proximity of the two proteins is required to demonstrate ternary complex formation.

If the EC_50_ value of a ternary binding experiment in a target engagement assay (%T_bound, ternary_ curve) coincides with *K*_T,2_ no other binding events other than the recruitment of CL to T is expected at equilibrium (Equation 3). In principle, with species C, T and L, three bindings with T are possible in equilibrium:

1. T + L ⟶ TL,
2. TL + C ⟶ TLC,
3. T + CL ⟶ CLT.

If the experimental conditions are such that contributions from the binding events of 1) and 2) are practically negligible, the measured EC_50_ of the %T_bound_ curve exclusively represents the binding event described by 3) and thus describes the equilibrium in Equation 3. Such conditions can be enforced by a high concentration of the stronger binding counter protein, in this example of chaperone C. Since we assume a weak affinity of L to T and a strong affinity of L to C, all or close to all ligand L is supposed to be bound to C in the chaperone–ligand complex CL if [C] is, for instance, 100-times higher than *K*_C,1_ (referred to as “saturation conditions”).

This situation is illustrated in the following example of a typical molecular glue of type I:

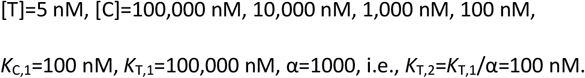

Derived parameters computed with the comprehensive mathematical model at equilibrium for [C]=10,000 nM and [L_tot_]=100 nM are: [TL]=2.48·10^−5^ nM, i.e., TL represents 0.0005% of T_tot_.^29^ When the excess concentration of C is lowered to 1000 nM, the concentration of [TL] is still only 0.002 nM, which corresponds to less than 0.005% of bound target T. For [C]=100 nM, TL increases to 0.05%, which is still negligible. The 1000-fold higher affinity of L for C relative to T and a 20-fold excess of C (100 nM vs. 5 nM) is sufficient to absorb almost all ligand into CL and suppress any significant concentration of TL. At [L_tot_]=100 nM, [CL] corresponds to 37.4 nM due to the 20-fold excess of C over T, the EC_50_ value of the corresponding %T_bound, ternary_ curve is 5000 nM, which is far away from *K*_T,2_=100 nM and the concentration of TL comprises about 2.5% of [T_tot_]=5 nM. This means that at such excess of the more strongly binding counter protein C, ternary complex formation at equilibrium comprises practically only the recruitment of CL to T with no significant concentration of TL. Consequently, the concentrations of T and CLT coincide at the inflection point of the %T_bound, ternary_ curve. This case is represented by Equation 3 when [T]=[CLT] and, hence, *K*_T,2_=[CL]. Moreover, if [C] is sufficiently high, all [L_tot_] is absorbed by C into CL and, hence, [CL] approaches [L_tot_], and consequently the EC_50_ of the %T_bound, ternary_ curve approximates the ternary *K*_d_ *K*_T,2_.

It is obvious that although a 20-fold excess of C is sufficient to suppress significant TL formation around the ternary *K*_d_ of *K*_T,2_=100 nM, this excess is insufficient to approximate [L_tot_] to [CL] and enable the estimation of *K*_T,2_ via the EC_50_ of the %T_bound, ternary_ curve. If the concentration of C is increased to a 200-fold excess over T, i.e., [C]=1000 nM, [CL] increases to 88 nM at [L_tot_]=100 nM. The EC_50_ of the %T_bound, ternary_ curve becomes 113 nM, which is still away from *K*_T,2_=100 nM. For [C]=10,000 nM, [CL] at [L_tot_]=100 nM corresponds to 96.6 nM. The EC_50_ of the %T_bound, ternary_ curve is 103.5 nM, which is close to the theoretically possible 102.5 nM (Equation 17 below).

It is of interest to study at what excess of C “all” ligand is absorbed by C into CL, i.e., at what excess of C L_tot_ (to which the EC_50_ refers to) corresponds to [CL]. If [C] is increased to an exorbitant 100,000 nM, [CL] approaches a value of 97.5 nM at [L_tot_]=100 nM but does not converge towards *K*_T,2_=100 nM. The remaining difference is 50% of [T]=2.5 nM. To achieve an EC_50_ of the %T_bound, ternary_ curve of 100 nM an [L_tot_] of 102.6 nM is required. This relationship is derived algebraically below. Intuitively, at [L_tot_]=100 nM the %T_bound, ternary_ curve reaches 50% of CLT formation if all [L_tot_] is bound into CL and CLT and [CL] cannot exceed 97.5 nM at this [L_tot_] regardless of [C]. Conversely, the EC_50_ value of the %T_bound, ternary_ curve cannot fall below 102.5 nM regardless of the excess of C over T (Fig.1).

**Fig. 1.**
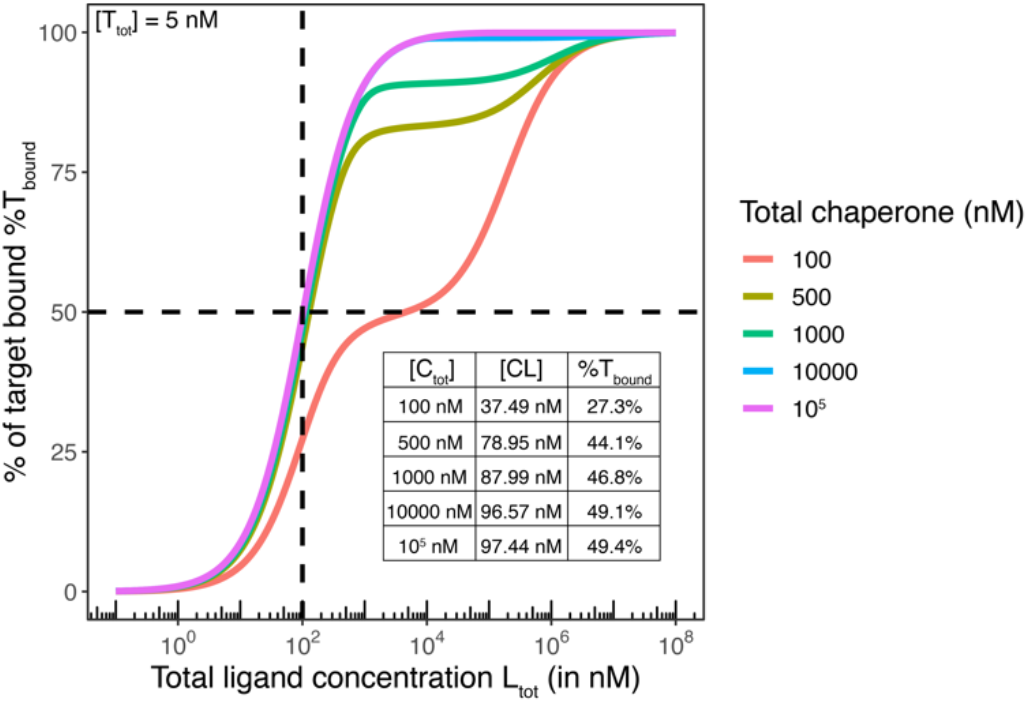
Saturation as a function of the chaperone concentration. For a typical molecular glue example with [T_tot_]=5 nM, *K*_C,1_=100 nM, *K*_T,1_=100,000 nM, *K*_T,2_=100 nM and α=1000, EC_50_ values of calculated %T_bound, ternary_ curves approach *K*_T,2_=100 nM with increasing concentrations of the chaperone (calculated EC_50_s for 4,600 nM is 102.5 nM). The EC_50_ of 102.5 nM at [C_tot_]=100,000 nM corresponds to the theoretically estimated bound. The remaining difference is 50% of [T_tot_] because at the inflection point – under saturation conditions – 50% of [T_tot_]=5 nM is complexed by CL into CLT. Another angle to look at saturation is the CL concentration. As the chaperone concentration increases, the concentration of free ligand L vanishes and the entire ligand concentration is bound into CL and CLT. In this example, saturation is only reached at [C_tot_] =100,000 nM when [CL] approaches the theoretical maximum of 97.5 nM. Calculations are performed with the previously published comprehensive mathematical model.^29^

In conclusion, [C] must be about 100 times the value of *K*_C,1_ for the EC_50_–[T_tot_]/2 of the %T_bound, ternary_ curve to coincide with *K*_T,2_ (see below). If these conditions are experimentally met, measured EC_50_ values derived from the %T_bound, ternary_ curve in a target engagement assay can interpreted as *K*_T,2_ and consequently be used to determine the cooperativity α without the use of a mathematical model.

##### The impact of the concentration of the monitored weaker binding protein

To understand the influence of the concentration of the monitored weaker binding protein and whether the EC_50_ value of the %T_bound, ternary_ curve and *K*_T,2_ coincide, it is necessary to relate the corresponding EC_50_ and *K*_d_ or L_tot_ and free L respectively to each other. We have seen in the previous section that under certain conditions, the ternary complex formation equilibria can be simplified such that only one path needs to be considered. If we focus on this case, the influence of the concentration of the monitored, weaker binding protein can be analysed as part of a binary equilibrium in which CL binds to T and forms CLT (Equation 3). Note that in Equation 3, the “ligand” that binds to T is CL and not L. For clarity, and because of the general nature of the topic to binary or ternary equilibria, the influence of the concentration of the monitored weaker binding protein on whether the EC_50_ value matches *K*_T,2_ or not is discussed below for binary binding of L to C, but the entire concept can be applied directly to any binary equilibrium, including that of Equation 3.

##### Concept of EC_50_ and *K*_d_ in binary binding equilibria

In a binary equilibrium, the dissociation constant (*K*_d_) refers to the free ligand concentration at which 50% of the protein is bound to the ligand while the half-maximal concentration EC_50_ refers to the corresponding total ligand concentration:

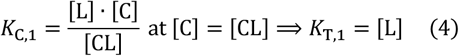

The *K*_d_ for protein C and ligand L (in a defined buffer and at a defined temperature) is a natural constant, i.e., it is independent of the experimental conditions and particularly of the selected protein C concentration. Since the determination of a *K*_d_ by measurement of the concentrations of three components of a binary equilibrium ([C], [CL] and [L]) requires non-trivial calibration of the analytical signals, scientists like to use, for practical reasons, EC_50_ values that refer to the required total ligand concentration L_tot_ at which 50% of protein binding is observed. EC_50_ values are well suited for the classification of compounds under the same assay conditions. However, it is mandatory for comparing compounds across assays to have the compound-specific and assay-independent *K*_d_ values.

To establish the relationship between the easily accessible EC_50_ and the *K*_d_, we start from the simple fact that the total concentration of protein C in a binary binding experiment is the sum of free and ligand-bound protein:

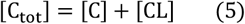

Assuming constant total protein, the half-maximal effect is reached at the total ligand concentration referring to 50% binding, the EC_50_ or [L_tot_]EC_50_:

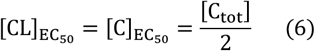

This inflection point is hence referring to the total ligand concentration at which bound and free C concentrations are balanced. Combining Equation 6 and the left-hand side of Equation 4, the identity of *K*_d_ and free ligand concentration at the inflection point is established:

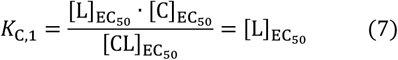

Using the identity for the total ligand at the inflection point, we get:

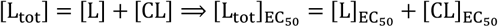

where [L_tot_]EC_50_is the measured EC_50_ value with respect to total ligand in the binary binding experiment. Substitution into Equation 7 yields the final result

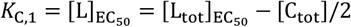

Alternatively, if we rearrange Equation 4 from above

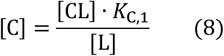

and substitute [C] in (8) with this definition, we obtain

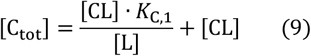

If we expand [CL] in (9) by [L] in numerator and denominator and obtain

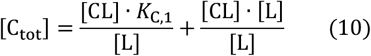

After combining the two summands, the equation becomes

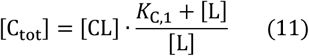

Rewriting (11) for [CL] gives

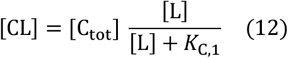

Equation (12) still contains [L], which is replaced in analogy to C_tot_ in equation (5):

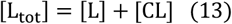

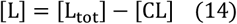

Once we substitute [L] into equation (9) with equation (11), we obtain

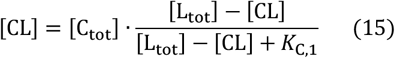

We now rearrange this equation in two steps such that *K*_C,1_ is placed on the left

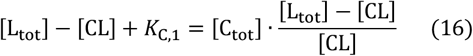

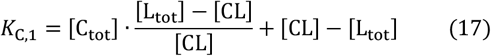

Equation (17) finally defines the dissociation constant independent of concentrations of free ligand and of free protein C but as a function of the known [L_tot_] and [C_tot_]. The only unknown concentration in Equation 14 is [CL], which is [C_tot_]/2 at the inflection point:

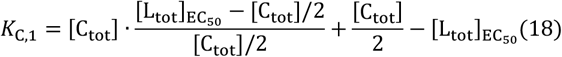

Equation 18 can be simplified to:

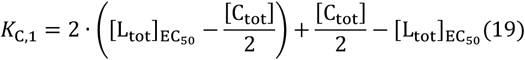

Or

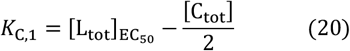

Equation (20) means that the *K*_d_ in a binary binding experiment can be obtained from the measured EC_50_ by subtracting half the protein concentration used. This explains the general finding that the EC_50_ is equal or close to the *K*_d_ value when the protein concentration is well below the *K*_d_ value but that the EC_50_ and the *K*_d_ differ significantly when the protein concentration is of the order of magnitude of the *K*_d_. For example, if the EC_50_=350 nM and the protein concentration in the corresponding binding experiment is [C_tot_]=500 nM, the *K*_d_ of this compound is 100 nM (350 nM−500/2 nM=100 nM). Intuitively, for a protein concentration around or greater than the *K*_d_ most of the compound is bound. Consequently, if in the same experimental environment a concentration of the free ligand [L] equals *K*_d_, i.e., when [CL] corresponds to [C], [L_tot_] needs to be well above *K*_d_, e.g., 350 nM in the above example (Fig. 2).

**Fig. 2.**
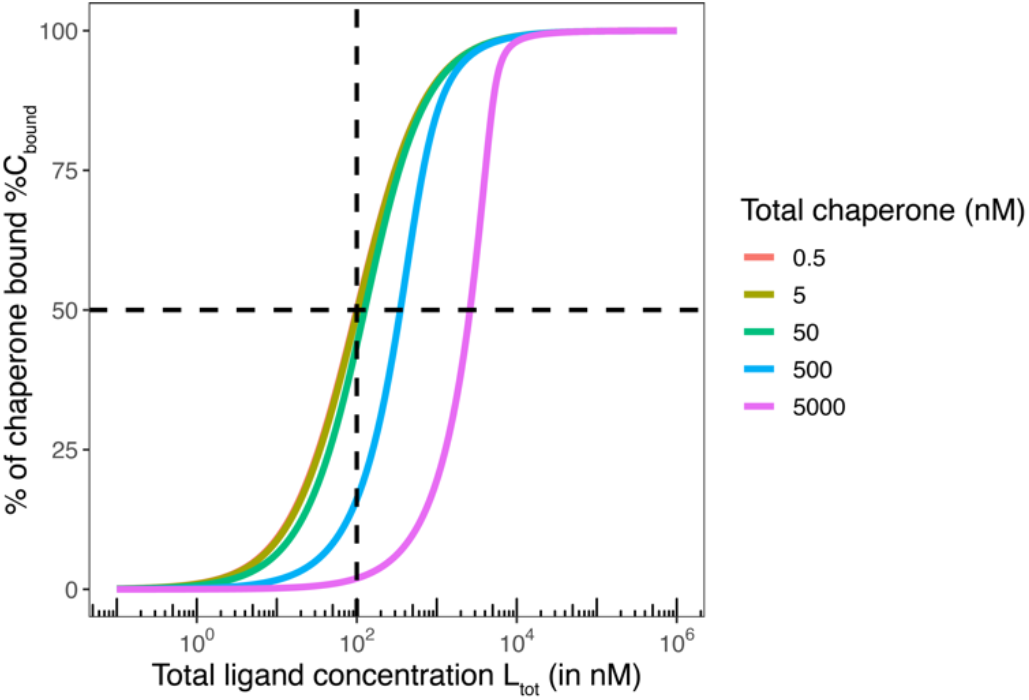
The monitored protein concentration influences the EC_50_ value of the %C_bound, binary_ curve. Protein concentrations above the *K*_d_ (here *K*_C,1_=100 nM) cause the EC_50_ and *K*_d_ to deviate from each other. For [C_tot_]=500 nM, the EC_50_ corresponds to 350 nM (blue curve), while for [C_tot_]=5 nM, the EC_50_ approaches 102.5 nM but cannot be lowered. Only at a protein concentration less or equal 200 the *K*_d_, the EC_50_ practically coincides with *K*_C,1_ (100.25 nM vs. 100 nM). These considerations also apply to the monitoring of ternary complex formation experiments under saturation conditions when the entire ligand concentration at the inflection point is bound into CLT and CL (or TL).

Equation 17 must now a) be applied to equation 3 and b) be combined with what was discussed in the previous section.

a. According to equation 17, the EC_50_ of a %T_bound_ curve with a concentration of the monitored protein T of 5 nM and a *K*_d_ of 100 nM cannot be closer to *K*_C,1_ than [L_tot_]EC_50_ =102.5 nM.
b. According to the relationship discussed above, a ternary complex-formation equilibrium can be treated as a binary equilibrium if the excess of the stronger binding protein C is about 20 times higher than that of the monitored weaker binding protein. If the excess of [C_tot_] over [T_tot_] is greater than 100-fold higher, [CL] coincides with [L_tot_]EC_50_, which means that the EC_50_ of the %T_bound, ternary_ curve approaches the limit of Equation 17.

To summarize, a 100-times higher [C_tot_] than *K*_C,1_ allows treating the ternary equilibrium as a binary system consisting only of CL and T as reactants. Moreover, it allows considering [L_tot_] as the concentration of the free ligand L that is CL, i.e., [L]=[CL], which means that the EC_50_ is the same as *K*_T,2_.

#### The influence of *K*_T,1_ on whether apparent and intrinsic cooperativity coincide

A last point of consideration is the definition of the “weaker binding protein”, i.e., at which value of *K*_T,1_ a significant TL formation starts to destroy the above discussed simplification into a binary system, despite the above-mentioned 100-fold excess of [C_tot_] over *K*_C,1_., This threshold represents then the boundary between molecular glues of type 1 and bi-functional compounds.

The validity of the above reduction of a ternary complex-forming equilibrium into a binary equilibrium is depending on the ratio of [C_tot_] and [T_tot_], as well as the ratio of the lower *K*_C,1_ and the higher *K*_T,1_. In the above-mentioned example with [T_tot_]=5 nM, [C_tot_]=10,000 nM, *K*_C,1_=100 nM and cooperativity α=1000, *K*_T,1_ can be as low as 10,000 nM until the difference between the EC_50_ of the %T_bound, ternary_ curve and *K*_T,2_ becomes significantly larger than the inherent and unavoidable difference, which is 50% of [T_tot_] or 2.5 nM. As discussed above, this is because the protein concentration of [T_tot_]=5 nM is too close to the corresponding *K*_T,2_ of 10 nM (=*K*_T,1_/α=10^4^ nM/1000). More specifically, the EC_50_ of the %T_bound, ternary_ curve is 12.6 nM, which is close to the theoretical 12.5 nM and the EC_50_ deviates already 25% from the *K*_d_. For *K*_T,1_>1000 nM, the relative difference falls below 25%.

Assuming the same parameters as above but with reduced α=100, *K*_T,2_ becomes 100 nM (=10^4^ nM/100), i.e., it deviates from [T_tot_] and the difference to the EC_50_ value of %T_bound, ternary_ is further reduced. This illustrates that a lower *K*_T,2_ only becomes relevant in combination with a high intrinsic cooperativity α and for *K*_T,2_ approaching the typical protein concentrations, which results in larger differences between the EC_50_ of the %T_bound, ternary_ curve and the underlying *K*_T,2_ (Fig. 3).

**Fig. 3.**
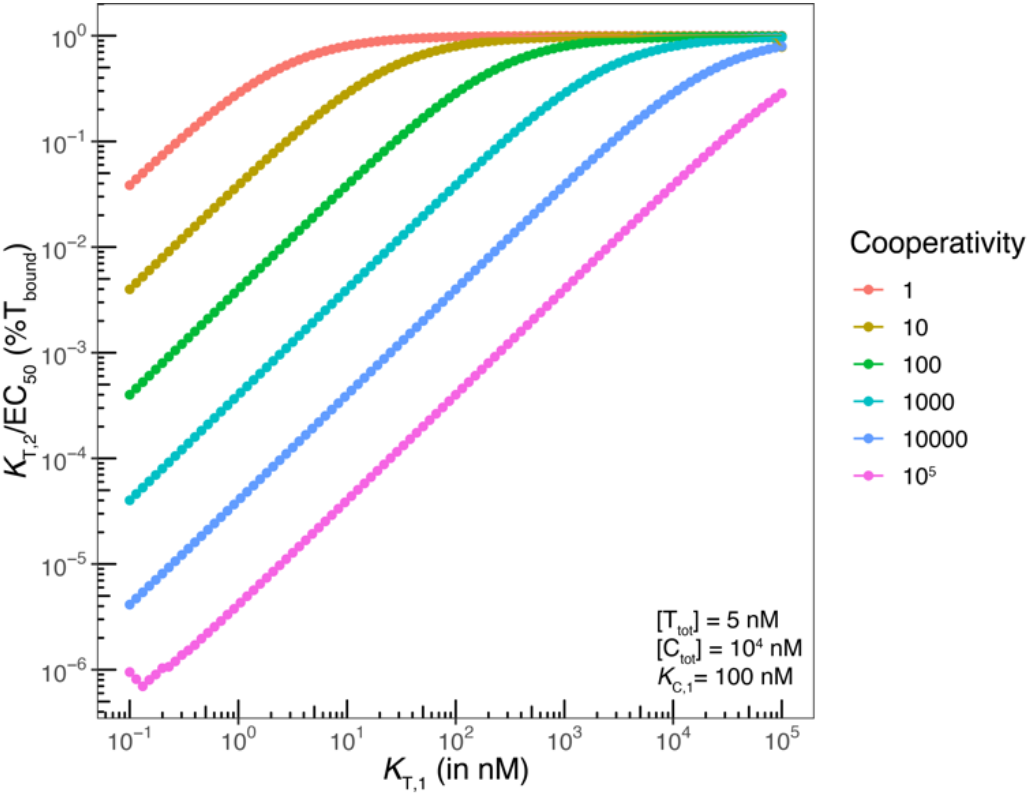
The deviation of the reduced pseudo two-body equilibrium with only one pathway and the three-body equilibrium comprising two pathways depends on the concentration of the relevant protein C. With the used [C_tot_]=10^4^ nM, the second pathway through TL becomes more relevant for lower *K*_T,2_=*K*_T,1_/α and the pseudo two-body equilibrium (“saturation conditions”) is affected. This leads to an increase in the ratio of *K*_T,2_ and EC_50_ of the %T_bound, ternary_ curve (y-axis) as *K*_T,2_ is close to or even below the concentration of the monitoring protein [T_tot_]=5 nM depending on the cooperativity α. In a non-cooperative system (α=1), saturation conditions are maintained down to a low *K*_T,1_≈10 nM (red curve). For higher cooperativity (α between 10 and 1000), significant differences between *K*_T,2_ and the EC_50_ of the %T_bound, ternary_ (brown, turquoise, blue curves) are observed for *K*_T,1_<10,000. For α>1000, *K*_T,1_ must exceed 100,000 nM to maintain saturation conditions (blue, cyan curves).

**Fig. 4.**
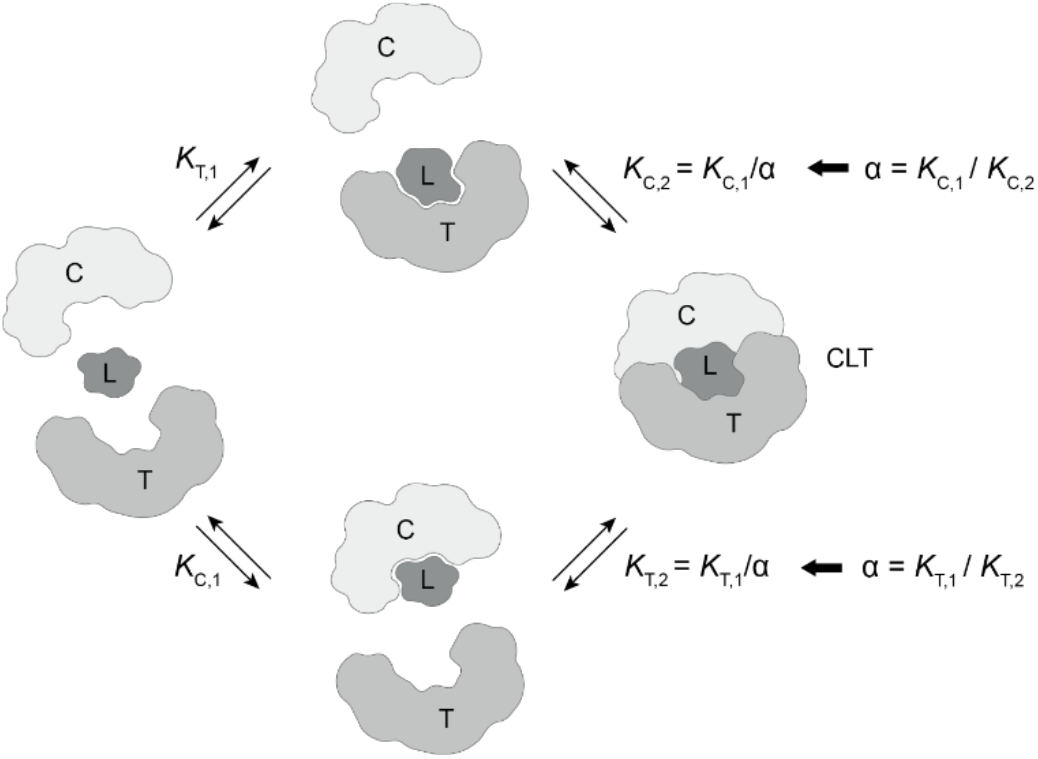
The formation of ternary complexes (CLT) is achieved through two different intermediate binary complexes (CL or TL). For type 1 molecular glues, L has a good to strong affinity to the chaperone protein C and a weak or very weak affinity to the chaperone protein T. A possible third pathway, the formation of binary chaperone– target complex followed by ligand binding (type 3 molecular glues) was excluded experimentally for the here presented examples.^**30**^ The intrinsic cooperativity α describes the factor of enhanced ligand binding of L to C when L is prebound to T (or to T when L is prebound to C). There is one unique cooperativity α that describes a given three-body system. The intrinsic cooperativity is, in analogy to the *K*_d_, a universal constant of the ternary complex and, therefore, independent of the experimental conditions. Maximizing the intrinsic cooperativity is a strong selection criterion in the drug development of molecular glues.

In summary, the difference between the EC_50_ value of the %T_bound, ternary_ curve and the underlying *K*_T,2_ becomes less than 2.5% with respect to *K*_T,2_ for [T_tot_] that is more than 20-times lower than *K*_T,2_ and [C_tot_] that is more than 100-times larger than *K*_C,1_. At the inflection point of the ternary complex-formation monitored by the target T, the concentration of the free target [T] is equal to the concentration of %T_bound_=[TL]+[CLT]. However, if all the free ligand is bound in [CL] (due to the excess of [C] and the weak affinity of L to T), [T_bound_]=[TL]+[CLT]≈[CLT] corresponds exclusively to [CLT], i.e., the inflection point of the %T_bound_ curve coincides with *K*_T,2_=[CL]≈[L_tot_] (see above). As soon as the affinity of L to T increases, the formation of the ternary complex is no longer the result of the binding of CL to T but also of the binding of TL to C. The same applies when the total concentration of C ([C_tot_]) becomes lower and approaches *K*_C,1_. In this case, the EC_50_ of the %T_bound, ternary_ curve (i.e., in the presence of the chaperone C) is not equal to *K*_T,2_ and the corresponding shift of the binary binding curve corresponds only to the apparent cooperativity and not the intrinsic cooperativity. In this case, both paths are significantly contributing and the intrinsic cooperativity can only be determined by a mathematical model, which requires input parameters from both binding assays with C and T as the monitoring proteins.

### Cooperativity studies on a small series of FKBP12:R,S-SLF-X:MAPRE1 ternary complexes

MAPRE1 is a plus-end tracking protein (+TIPs) that regulates microtubule (MT) behavior and interactions between MTs and other intracellular structures during mitosis._31_ It was identified as the recruitment complement of FKBP12 by a protein array screening of 50 macrocyclic FKBP12 ligands against 2500 randomly selected proteins. The ternary complex of FKBP12:**R,S-SLF-1a**:MAPRE1 was characterized previously.^30^ The protein NMR and X-ray structure of the ternary complex reveal multiple interactions between **R,S-SLF-1a** and MAPRE1 as well as interactions between FKBP12 and MAPRE1, both of which only occur in the ternary complex. The increased binding of **R,S-SLF-1a** and related compounds to FKBP12 by the presence of MAPRE1 or to MARPE1 by the presence of FKBP12 enables the retrieval of the intrinsic cooperativity α. Due to the distinct binary affinities of the studied molecular glues, we expect different apparent cooperativities but the same intrinsic cooperativity α regardless of whether monitoring is through MAPRE1 or FKBP12.

### Compound selection for cooperativity from %A_max_ values based on TR-FRET proximity assay

TR-FRET proximity assays quantify the compound-mediated induced proximity by measuring the distance-dependent energy transfer between a donor- and an acceptor-tagged protein. It is noteworthy that ternary complex-forming compounds do not necessarily have to exhibit intrinsic cooperativity. Even in the absence of cooperativity (α=1), ternary complexes can be formed based on the independent affinity of such compounds to either of the two proteins. While a TR-FRET proximity assay can monitor the formation of a non-cooperative ternary complex formation, i.e., *K*_C,1_/*K*_C,2_=*K*_T,1_/*K*_T,2_=1, a target engagement assay cannot as it only reports increased ligand binding caused by interactions that occur exclusively in the ternary complex (α>1). However, if no ternary complex is formed at all, there is also no compound-dependent increase in the %A_max_ value of the TR-FRET assay. For this reason, we focus on those compounds from our previous publication^30^ that exhibited the highest %A_max_ values in the TR-FRET proximity assay when evaluating cooperativity with a target engagement assay (Table 1).

**Table 1.**
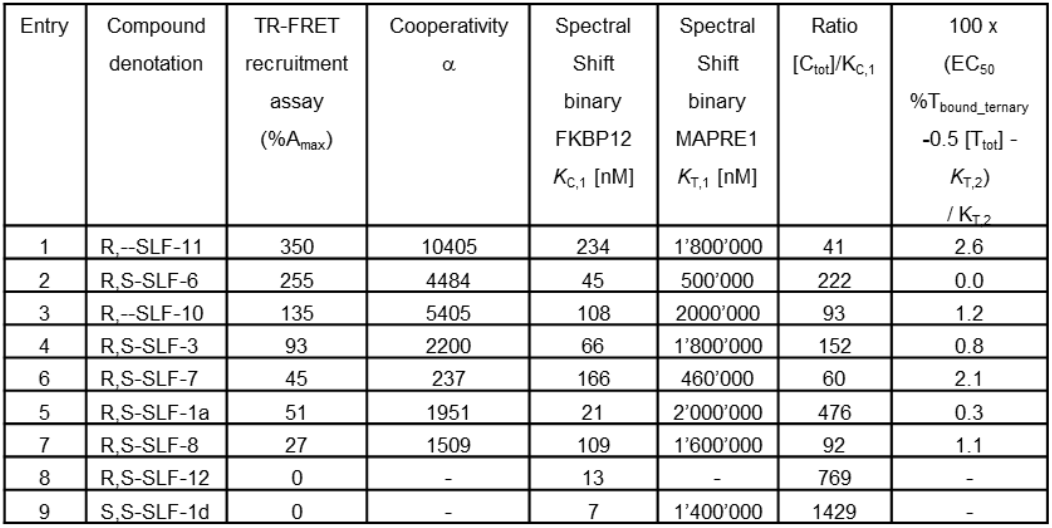
Column 3 shows the %A_max_ values from the TR-FRET proximity assay, reported earlier.^**30**^ Column 4 shows the cooperativity α values calculated from measured *K*_C,1_, *K*_T,1_ and the EC_50_ value of the %T_bound, ternary_ curve discussed in this work. Column 5 and 6 show *K*_C,1_ and *K*_T,1_, respectively, obtained through the spectral shift target engagement assays. Column 7 shows the ratio from [C_tot_] and *K*_C,1_. Column 8 shows in percentages the difference between the EC_50_ of the %T_bound, ternary_ curve and the corresponding *K*_T,2_.

All TR-FRET-active compounds from R,S-SLF-1a to R,**-**-SLF-11 have the R-configuration at the α-position of the 2-amido-4-methylene-pipecolinic moiety and the S-configuration at the α-position of the second amino acid of the recruitment loop (if there is a chiral center). The compounds S,S-SLF-1d and S,S-SLF-12 are negative controls that bind strongly to FKBP12 but lack any %A_max_ increase, i.e., do not recruit MAPRE1 (%A_max_=0%) (Fig. 5).

**Fig. 5.**
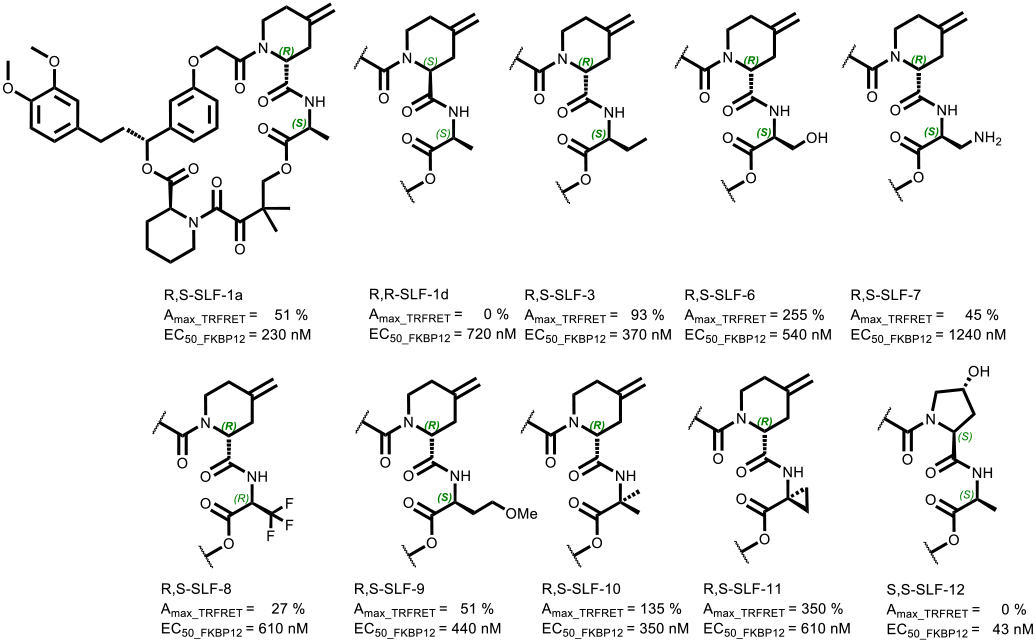
Ten selected compounds from our previous publication on MAPRE1 for cooperativity studies.^**30**^ Eight of them show significant %A_max_ values in the TR-FRET proximity assay indicating ternary complex-formation. Two compounds (**S,S-SLF-1d and S,S-SLF-12**) were selected as negative controls. **S,S-SLF-1d** differs only in the stereochemistry at the α-position of the 2-amido-4-methylene-pipecolinic moiety compared **to R,S-SLF-1a**.

#### Binding studies on MAPRE1 and FKBP12 using a spectral shift target engagement assay

As discussed in the previous section, the study of the apparent cooperativity consists of two steps: the binding characterization of a ligand to a target protein T in the absence and in the presence of a counter protein. The counter protein will (in case there is intrinsic cooperativity) lead to an increased affinity of the ligand for T (or lower *K*_T,2_) relative to the ligand alone (*K*_T,1_). Importantly, the requirement that both experiments are identical (except for the presence of the counter protein) allows labelling of only the protein by which the binding event is monitored. An alternative to this assay without protein labelling would be a competitive binding assay which measures changes in fluorescence polarization by a fluorescent tracer–ligand substitution whose fluorescence signal changes significantly when it transitions from a bound to an unbound state. However, a competitive binding assay requires chemical matter that binds to the target. Since there is no reason to expect strong binding of the FKBP12:**R,S-SLF-X**:MAPRE1-forming compounds to MAPRE1 alone, a prerequisite to developing the competitive binding assay was a reporter system. Such system monitors only subtle changes on the target but has sufficient sensitivity to measure very weak binding to enable monitoring of the apparent cooperativity by the weaker binding protein. Compared to a competitive binding assay, the monitoring of binding by structural changes on the protein is independent of the binding site.

Our method of choice is the spectral shift method, which monitors binding events on the target isothermally and in equilibrium via an environment-sensitive near-infrared fluorophore.^32^ In the case of target engagement either the proximity of the ligand towards the dye or structural changes occurring in the target as a result of binding lead to a change in the chemical environment of the fluorophore. Those changes affect the physiochemical properties of the reporter fluorophore and translate into either blue or red shifts (or peak broadening) of emission peaks upon ligand binding, which can be sensitively monitored by simultaneously recording the fluorescence at two wavelengths (i.e., at 650 nm and 670 nm). Calculating the ratio between the two fluorescence channels then allows to monitor subtle emission shifts that occur as a result of target engagement. Affinity measurements can then be performed by recording the spectral shift as a function of ligand concentration.

In the following section, the binding of each of the ten selected compounds through MAPRE1 and FKBP12 was studied and the resulting apparent and intrinsic cooperativities were compared in an A_max_-ranked order.

#### R,S-SLF-11: selected compound with highest %A_max_ value of 350%

##### Binding study through MAPRE1

MAPRE1 was labelled via lysines using an NHS-conjugated dye. The final concentration of MAPRE1_NHS_ (referring to [T_tot_]) was 5 nM. Compounds could be titrated up to a concentration of 200 μM (referring to [L_tot_]). Titration of **R,--SLF-11** up to [L_tot_]=200 μM showed a very clear well-characterized onset of the %T_bound, binary_ curve, which by interpolation, resulted in an estimated EC_50_ of 1.8 mM. For well-defined minimal response values of the dose–response curve fitting of standard models resulted in reasonable approximations for EC_50_ and derived *K*_d_=EC_50_–[T_tot_]/2. However more accuracy is only achieved at higher ligand concentrations – ideally at full target saturation. However, this is often experimentally infeasible due to assay conditions (i.e., the requirement of maintaining a certain DMSO concentration or avoiding solubility issues of the ligand). Although the determined *K*_T,1_ values of binding to MAPRE1 are only approximations, their relative ranking reflects the underlying differences in affinity of the binary complexes. Performing these experiments in duplicates confirmed the high signal-to-noise ratio, i.e., there was no significant deviation between the two curves, indicating a high accuracy of the interpolated EC_50_ values. Due to the low [T_tot_]=5 nM, this value coincides with *K*_T,1_ (1.8 mM–0.005 mM/2≈1.8 mM). Since FKBP12 is a well-behaved protein, an excess of [C_tot_]=10 μM could be used for *K*_d_ characterization resulting in an EC_50_ value of the %T_bound, ternary_ curve of [L_tot_]=180 nM and an apparent cooperativity=10,000. Since [C_tot_]=10 μM is only 41 times the *K*_C,1_=234 nM instead of the recommended factor of 100 (Fig. 6 left), the EC_50_ value of 180 nM of the measured %T_bound, ternary_ curve does not exactly match the value from the general mathematical model. Iteration until the intrinsic cooperativity α matches the measured EC_50_ value of [L_tot_]=180 nM (at [C_tot_]=10,000 nM, [T_tot_]=5 nM, *K*_C,1_=234 nM, *K*_T,1_=1,800,000 nM) results in an intrinsic cooperativity of α=10,405 (Fig. 6 right). At [L_tot_]=180 nM, the calculated corresponding “ligand” concentration of [CL] is 173 nM, which is below the upper maximum of [CL]=180 nM–2.5 nM=177.5 nM. This means that with a [C_tot_] of “only” 10 μM, [L_tot_]=180 nM does not achieve the maximum possible concentration of [CL]=177.5 nM unlike an excess of [C_tot_] of 100 times *K*_C,1_ (or higher). Consequently, *K*_T,2_ corresponds to 173 nM and not 177.5 nM. This deviation of 2.6% between the measured EC_50_ of the %T_bound, ternary_ curve and the underlying *K*_T,2_ represents the difference of the measured apparent cooperativity of 10,000 and the corresponding underlying intrinsic cooperativity α=10,405 (Fig. 6 right).

**Fig. 6.**
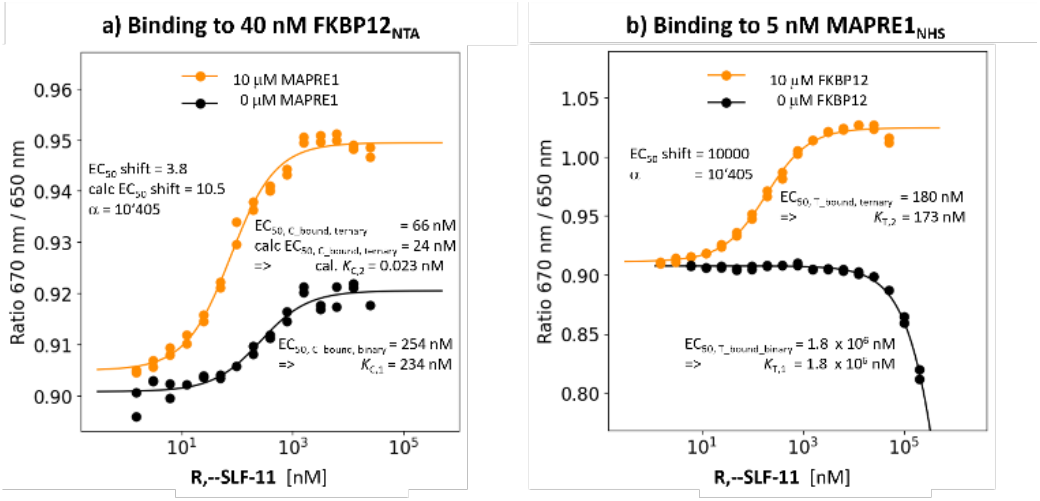
Binding data with experimentally derived and calculated cooperativity and *K*_d_s for R,--SLF-11. Right: The interpolated EC_50_ of the %T_bound, binary_ curve for binary binding to MAPRE1 is 1.8 mM. Due to the low [T_tot_]=5 nM, this EC_50_ was interpreted directly as *K*_T,1_. The presence of 10 µM FKBP12 resulted in a shift of the EC_50_ of the %T_bound. Ternary_ curve to 180 nM, which corresponds to an apparent cooperativity of 10,000. Left: EC_50_ of 254 nM of the %C_bound, binary_ curve corrected by 50% of [C_tot_]=40 nM to *K*_C,1_=234 nM. Using the full model with [C_tot_]=10,000 nM, [T_tot_]=5 nM, *K*_C,1_=234 nM, *K*_T,1_=1,800,000 nM and EC_50_ of 180 nM of the %T_bound, binary_ curve results in the cooperativity α=10,405 and *K*_T,2_=173 nM. Running the same experiment in the opposite direction, i.e., via the other binding protein, with the EC_50_=66 nM of the %C_bound, ternary_ curve (instead of 24 nM) results in an apparent cooperativity of 3.8 relative to the calculated 10.5 using α=10,405 from the EC_50_ of %T_bound, ternary_ curve. The trend to underestimating EC_50_-shifts by factor 2 to 3 when determining cooperativity through FKBP12 is constant across all compounds. Possible reasons are discussed in the main text. An α=10,405 results in *K*_C,2_=0.023 nM, which is below [C_tot_]=5 nM and indicates that the assay is bottoming out in for FKBP12.

##### Binding studies through FKBP12

To monitor the binding of **R,--SLF-11** by the stronger binding FKBP12 in the absence and in the presence of 10 μM MAPRE1 via %C_bound, binary_ and %C_bound, ternary_, respectively, the concentration of reversibly NTA-labelled FKBP12 was limited to [C_tot_]=40 nM. The EC_50_ value of the %C_bound, binary_ curve was 254 nM compared to 230 nM in the TR-FRET competitive binding assay from our previous work. This results in *K*_C,1_=234 nM using Equation 20. Repeating this experiment in the presence of [T_tot_]=10 μM of MAPRE1 resulted in an EC_50_ of the %C_bound, ternary_ curve of 66 nM corresponding to a relative shift or apparent cooperativity of 254 nM/66 nM=3.8. In contrast to the experimental setup in the previous two binding experiments with monitoring through the weaker binding protein MAPRE1, where the applied protein concentrations were optimally chosen, this is not the case for the binding experiment through the stronger binder FKBP12. First of all, the applied excess of T with [T_tot_]=10 μM is roughly 200-fold off of the proposed 100 times *K*_T,1_ threshold, which would correspond to 1.8 mM. Only under such excess conditions of T, the equilibrium of forming TL under the constraint that all ligand L is bound to T can be reached to ensure that [L_tot_] approaches the “ligand” concentration [TL]. Moreover, the concentration of the labelled protein FKBP12_NTA_ with [C_tot_]=40 nM is too close to the corresponding *K*_C,1_=234 nM resulting in a high offset of 20 nM=[C_tot_]/2 of the EC_50_ value of the %C_bound, ternary_ curve and the corresponding *K*_C,2_. If *K*_C,2_ is sufficiently high, the 20 nM deviation to correct for the protein concentration could be neglected. However, the resulting calculated value for *K*_C,2_ is 0.023 nM accounting for the cooperativity α=10,405 and *K*_C,1_=234 nM as derived by the MAPRE1 experiment. This value is far below the applied protein concentration of [C_tot_]=40 nM and makes it infeasible to estimate *K*_C,2_ from the EC_50_ of the %C_bound, ternary_ curve. Consequently, the EC_50_ shifts, meaning the apparent cooperativity, from monitoring the binding by the stronger binding protein C are diverging from the underlying intrinsic cooperativity α. Using the binary and ternary binding data to MAPRE1 with cooperativity α=10,405 and *K*_T,1_=1.8 mM and the binary binding to FKBP12 with *K*_C,1_=234 nM, a resulting EC_50_=24.2 nM is estimated for the %C_bound, ternary_ curve relative to the experimentally determined 66 nM. The deviation of calculated (254 nM/24.2 nM=10.5) and measured apparent cooperativity (254 nM/66 nM=3.8) can be explained by a negative cooperativity induced by the His6-NTA tag of FKBP12. Studies to lower [C_tot_] and identify a better position for the spectral shift tag on FKBP12 are ongoing (Fig. 6 left).

#### R,S-SLF-6: second highest %A_max_ value of 255%

##### Binding study through MAPRE1

The compound with the second highest A_max_ of 255%, **R,S-SLF-6**, is found to have an EC_50_ value of the %T_bound, binary_ curve of 0.5 mM. The EC_50_ value of the %T_bound, ternary_ curve was shifted to 114 nM in the same binding experiment but in the presence of 10 μM FKBP12, i.e., [C_tot_] is 222 times the *K*_C,1_. From the mathematical model a “ligand” concentration of [CL]=111.5 nM at [L_tot_]=114 nM is found. This means that any applied ligand L_tot_ that is not bound into CLT is bound to CL, i.e., [CL]≈EC_50_– [T_tot_]/2≈*K*_T,2_≈111.5 nM. Consequently, the EC_50_ value of the %T_bound, ternary_ curve matches *K*_T,2_ exactly for the given [T_tot_]=5 nM (0% deviation). As a result, the apparent cooperativity of 500,000 nM/114 nM=4386 coincides with the intrinsic cooperativity α of 500,000 nM/111.5 nM=4484 and the systematic difference due to the protein concentration becomes neglectable (Fig. 7 right).

**Fig. 7.**
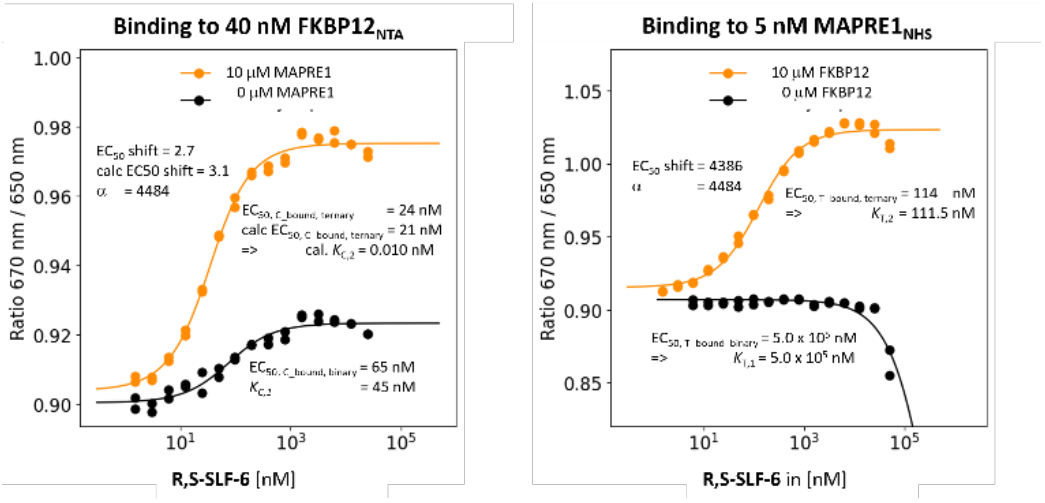
Binding and calculated data for R,S-SLF-6. For explanation of details refer to Fig. 6.

##### Binding study through FKBP12

The obtained results from the above binding studies via the alternative protein can be used to validate the proposed concept of retrieving the intrinsic cooperativity α from only one binding assay. For **R,S-SLF-6**, the EC_50_ values shift in the binary to ternary setup from 65 nM (i.e., *K*_C,1_=45 nM) to 24 nM. With a cooperativity α=4484, obtained via MAPRE1 binding, the EC_50_ value of the calculated %C_bound, ternary_ curve resulted in 21 nM, which is an almost perfect match and confirms the independently obtained values of the MAPRE1 binding experiment. The calculated “ternary” *K*_d_ *K*_C,2_=*K*_C,1_/α=45 nM/4,484=0.01 nM indicates that the EC_50_ value of the %C_bound, ternary_ curve of 21 nM is much greater than *K*_C,2_ resulting in distinct apparent and intrinsic cooperativity values which highlights that the information of the stronger binding protein alone is insufficient to characterize the intrinsic cooperativity α without using a comprehensive mathematical model (Fig. 7 left).

#### R,--SLF-10: third highest %A_max_ value of 135%

##### Binding study through MAPRE1 R,--SLF-10

the compound with third highest %A_max_ of 135%, has an interpolated EC_50_ value of the %T_bound, binary_ curve of only 2.0 mM and therefore with an even weaker potency than **R,S-SLF-11**. The same binding experiment but in presence of 10 μM FKBP12, i.e., [C_tot_]=93·*K*_C,1_, resulted in an EC_50_ shift of the %T_bound, ternary_ curve to [L_tot_]=377 nM. The “ligand” concentration from the mathematical model results in [CL]=370 nM at [L_tot_]=377 nM. This is a deviation of 1.2% from the theoretical upper bound of 377 nM–2.5 nM=374.5 nM. This also implies that *K*_T,2_ is not 374.5 nM as hinted by the EC_50_ value but 370 nM. Consequently, the apparent cooperativity of 2,000,000 nM/377 nM =5305 approaches the intrinsic cooperativity α=2,000,000 nM/370 nM=5405 (Fig. 8 right).

**Fig. 8.**
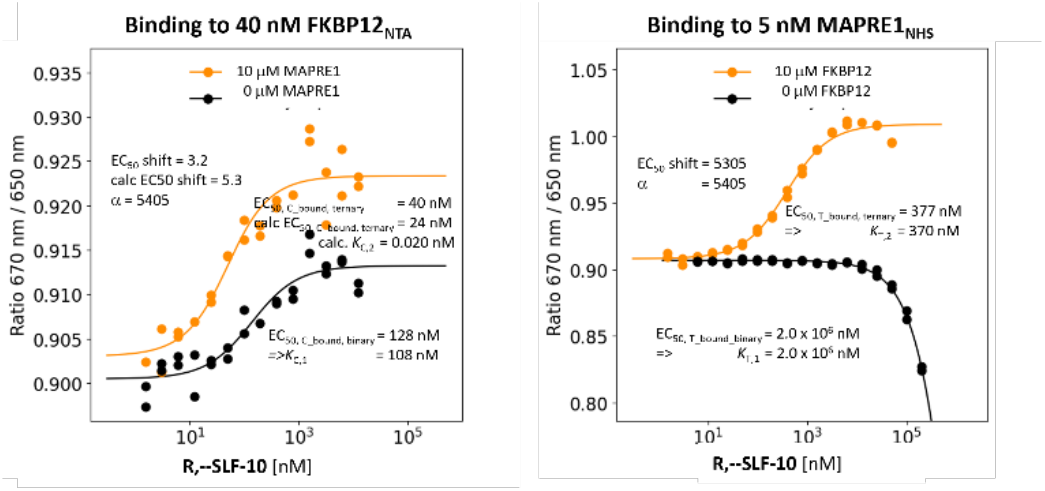
Binding and calculated data for R,--SLF-10. For explanation of details refer to Fig. 6.

##### Binding study through FKBP12 For R,--SLF-10

the EC_50_ of the %C_bound, binary_ curve shifted from 128 nM (i.e, *K*_C,1_=108 nM) to the EC_50_ of the %C_bound, ternary_ curve of 40 nM resulting in an apparent cooperativity of 3.2. With the cooperativity α=5405 from the MAPRE1 binding experiments the EC_50_ of the %C_bound, ternary_ curve is 24 nM with an apparent cooperativity of 5.3. The calculated “ternary” *K*_d_ becomes *K*_C,2_=*K*_C,1_/α=108 nM/5405=0.024 nM and it shows that the EC_50_ value of the %C_bound, ternary_ curve of 40 nM is far from *K*_C,2_=0.024 nM (Fig. 8 left).

#### R,S-SLF-3: %A_max_ value of 93%

##### Binding study through MAPRE1 For R,S-SLF-3

the compound with the fourth highest %A_max_ of 93% an EC_50_ value of the %T_bound, binary_ curve 1.8 mM was interpolated. The same binding experiment but in the presence of 10 μM FKBP12, i.e., [C_tot_]=152·*K*_C,1_, resulted in a shift of the EC_50_ value of the %T_bound, ternary_ curve to [L_tot_]=827 nM. The calculated model-based “ligand” concentration is [CL]=818 nM at [L_tot_]=827 nM. This is a rather small deviation of 0.8% from the true value of 827–2.5 nM=824.5 nM. This means that *K*_T,2_ is not 824.5 nM when directly reading off the EC_50_ but rather 818 nM. Moreover, the apparent cooperativity of 1,800,000 nM/823 nM=2177 approaches the intrinsic cooperativity α of 1,800,000 nM/818 nM=2200 (Fig. 9 right).

**Fig. 9.**
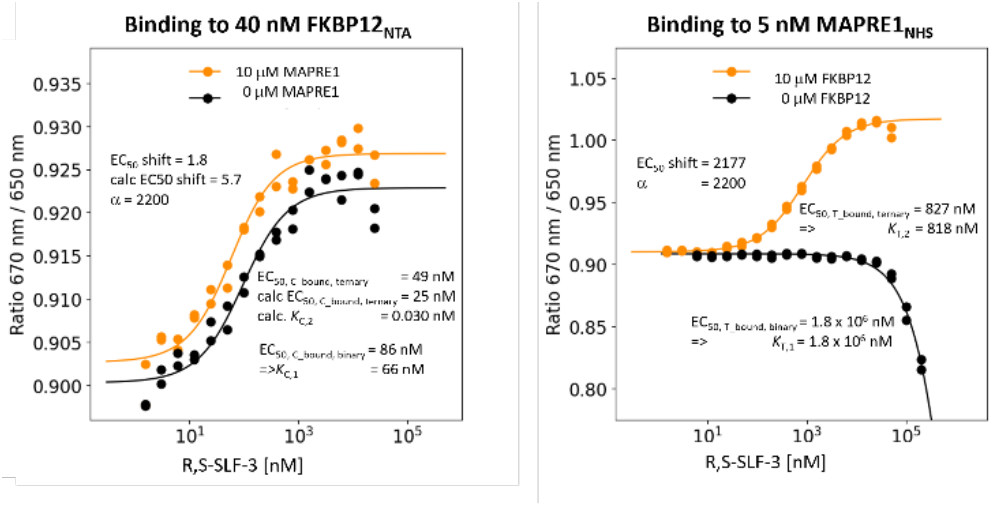
Binding and calculated data for R,S-SLF-3. For explanation of details refer to Fig. 6.

##### Binding study through FKBP12 For R,S-SLF-3

the EC_50_ of the %C_bound, binary_ curve shifted from 86 nM (then *K*_C,1_=66 nM) to a measured EC_50_ of the %C_bound, ternary_ curve of 49 nM resulting in an apparent cooperativity of 1.8 relative to a calculated apparent cooperativity of 5.7. The same value estimated with the cooperativity α of 2200 from the MAPRE1 binding experiment gave an EC_50_ of the interpolated %C_bound, ternary_ curve of 25 nM, which deviates from the measured value by factor of only factor 2 (5.3/1.8=1.9). This independently confirms the accuracy of the measurement through the weak MAPRE1 binding. The calculated “ternary” *K*_d_ of *K*_C,2_=*K*_C,1_/α=66 nM/2200=0.030 nM shows that the EC_50_ value of the %C_bound, ternary_ curve is far from *K*_C,2_, i.e., the monitoring of binding through the stronger binding protein does not allow the direct determination of *K*_C,2_ and consequently the intrinsic cooperativity α cannot be determined without a mathematical model describing all non-saturated conditions (Fig. 9 left).

#### R,S-SLF-7: absolute %A_max_ value of 45%

Due to a aminomethyl instead of the hydroxymethyl moiety (as **R,S-SLF-6**) at the at the second residue or the recruitment loop, **R,S-SLF-7**’s %A_max_ value drops from 255% to only 45%.

##### Binding study through MAPRE1

The EC_50_ value of the %T_bound, binary_ curve of **R,S-SLF-7** was interpolated to 0.46 mM, which is slightly lower than for the previous compounds. It has to be noted that R,S-SLF-7 showed precipitation at high concentrations. Therefore, measurements from the highest concentration samples had to be removed, which impaired the determination of the interpolation curve. Moreover, the signal is much smaller compared to the other studies ligands. In the same binding experiment but in the presence of 10 μM FKBP12 (i.e., [C_tot_]=60·*K*_C,1_) the EC_50_ of the %T_bound, ternary_ curve shifted to [L_tot_]=1985 nM. The model-based estimate is [CL]=1941 nM for the “ligand” at [L_tot_]=1985 nM. This is a deviation from the theoretically found true value of 1985 nM–2.5 nM=1982.5 nM. The *K*_T,2_ is 2% below the EC_50_ value of the %T_bound, ternary_ curve. The apparent cooperativity of 460,000 nM/1985 nM=232 is still below the cooperativity α of 460,000 nM/1941 =237 (Fig. 10 right).

**Fig. 10.**
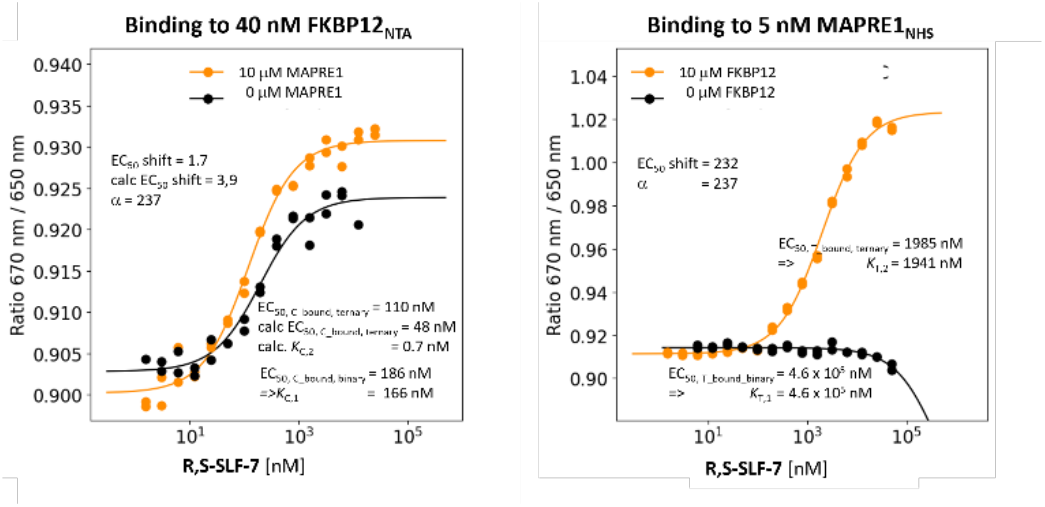
Binding and calculated data for R,S-SLF-7. For explanation of details refer to Fehler! Verweisquelle konnte nicht gefunden werden.

##### Binding study through FKBP12

The EC_50_ value for **R,S-SLF-7** shifted from 186 nM in the %C_bound, binary_ curve (resulting in *K*_C,1_= 166 nM) to a measured EC_50_ of the %C_bound, ternary_ curve of 110 nM, so the apparent cooperativity becomes 1.7. Using the cooperativity α=237 from MAPRE1 binding resulted in an EC_50_ value for the calculated %C_bound, ternary_ curve of 48 nM, which corresponds to a calculated apparent cooperativity of 3.9. The calculated “ternary” *K*_d_ of *K*_C,2_= *K*_C,1_/α=166 nM/237=0.7 nM shows that the EC_50_ value of the %C_bound, ternary_ curve is again very distinct from *K*_C,2_, i.e., the direct determination of *K*_C,2_ by monitoring the binding events corresponding to the stronger binding protein does not enable the direct identification of the cooperativity α without using a mathematical model (Fig. 10 left).

#### R,S-SLF-1a: absolute %A_max_ of 51%

**R,S-SLF-1a** represents the only hit compound from a protein array screening against 2500 proteins. Its %A_max_=51% in the TR-FRET proximity assay was the starting point that ultimately led to the identification of a series of glues with %A_max_ values of up to 350%, i.e., a 7-fold improvement with respect to %A_max_.

##### Binding study through MAPRE1

It was known from previous NMR studies that **R,S-SLF-1a** has a weak (greater than 250 μM) but measurable affinity for MAPRE1. This was confirmed by the binding study using the spectral shift method. The EC_50_ value of the %T_bound, binary_ curve of **R,S-SLF-1a** was estimated as 2.0 mM. The same binding experiment, but in the presence of 10 μM FKBP12, i.e., [C_tot_]=476·[*K*_C,1_], shifts the EC_50_ of the %T_bound, ternary_ curve to [L_tot_]=1030 nM. The model-based estimate of the “ligand” concentration yields [CL]=1025 nM at [L_tot_]=1030 nM. The *K*_T,2_=1025 nM deviates from the true value of 1030 nM–2.5 nM=1027.5 nM by 0.24%. **R,S-SLF-1a**’s apparent cooperativity of 2,000,000 nM/1030 nM=1942 is in great agreement with the cooperativity α=2,000,000 nM/1025 nM=1951) within the experimental error (Fig. 11 right).

**Fig. 11.**
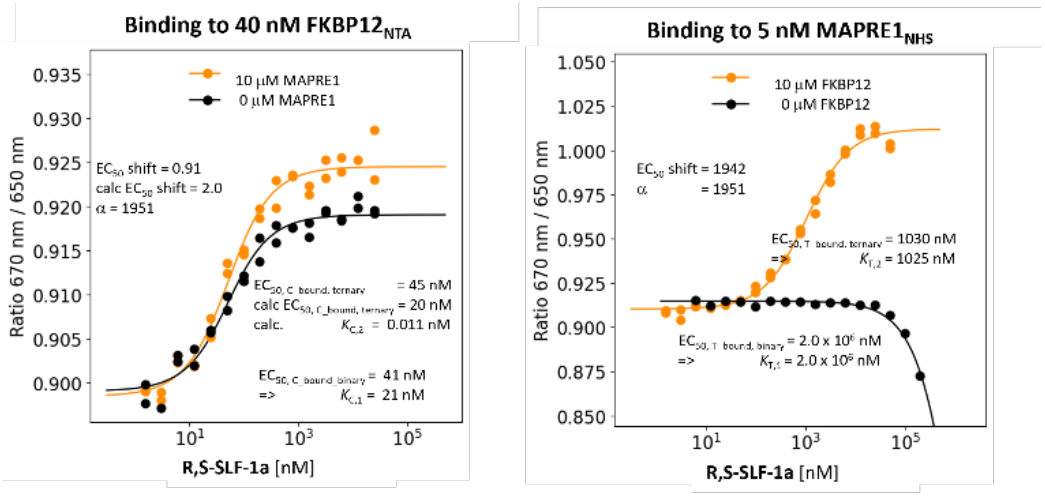
Binding and calculated data for R,S-SLF-1a. For explanation of details refer to Fig. 6.

**Binding study through FKBP12 R,S-SLF-1a**’sEC_50_ value of the %C_bound, binary_ curve of 41 nM (i.e., *K*_C,1_=21 nM) showed no significant shift in the EC_50_ of the %C_bound, ternary_ curve of 45 nM resulting in an apparent cooperativity of 0.9. Estimation using the cooperativity α=1951 from the MAPRE1 binding resulted in an EC_50_ for the calculated %C_bound, ternary_ curve of 22 nM, which is lower by factor 2 than the measured value. This result shows that the combination of a low *K*_C,1_ and a high [C_tot_] reaches the quantification limit of the assay. It is not possible for **R,S-SLF-1a** to detect an apparent cooperativity by monitoring the ternary binding formation by the stronger binding protein although there is significant underlying intrinsic cooperativity α. In this case, only the binding study by the weaker binding protein revealed the significant underlying cooperativity α. It is insightful to compare to **R,S-SLF-6** with an advantageous *K*_C,1_ of 45 nM, which is only 2.1-times higher than that of R,S-SLF-1a (21 nM). However, it is not the relative difference between the two values that is of relevance but the absolute value. The *K*_C,1_ of **R,S-SLF-1a** is exactly at the lower limit of the assay, which is *K*_C,2_+[C_tot_]/2=21 nM/1951+20 nM=0.011 nM+20 nM=20.011 nM. In other words, EC_50_s of the %C_bound, ternary_ curve below a threshold of 20 nM cannot be detected. If the EC_50_ of the %C_bound, binary_ curve is already at 20 nM, no shift is to be expected, which is seen in the experiment (Fig. 11 left).

#### R,S-SLF-8: %A_max_ of 27%

R,S-SLF-8 is the selected compound with the lowest %A_max_ value of 27% in the TR-FRET proximity assay.

##### Binding study through MAPRE1

The EC_50_ value of the T_bound, binary_ curve of **R,S-SLF-8** is interpolated to 1.6 mM. The same binding experiment but in the presence of 10 μM FKBP12, i.e., [C_tot_]=92·[*K*_C,1_], results in shifts of the EC_50_ of the %T_bound, ternary_ curve to [L_tot_]=1074 nM. The model-based concentration of the recruited “ligand” is [CL]=1060 nM at [L_tot_]=1074 nM. A *K*_T,2_=1060 nM means a deviation of 1.1% from the true value, which is 1074 nM–2.5 nM=1071.5 nM. This means that the apparent cooperativity of 1,600,000 nM/1074 nM=1490 for **R,S-SLF-8** matches the intrinsic cooperativity α =1,600,000 nM/1060 nM=1509 with a deviation that could be explained by the incomplete saturation (Fig. 12 right).

**Fig. 12.**
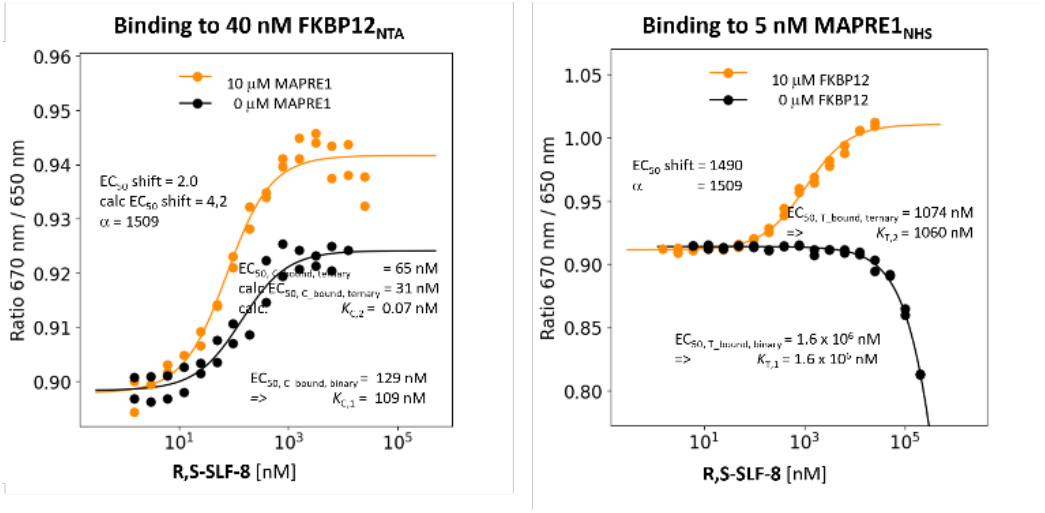
Binding and calculated data for R,S-SLF-8. For explanation of details refer to Fig 6.

##### Binding study through FKBP12

The EC_50_ of the %C_bound, binary_ curve of **R,S-SLF-8** is 129 nM (i.e., *K*_C,1_=109 nM) shifted to a measured EC_50_ of the %C_bound, ternary_ curve of 65 nM resulting in an apparent cooperativity of 2.0. Recalculating this with the intrinsic cooperativity α=1509 obtained through MAPRE1 binding gave an EC_50_ of the calculated %C_bound, ternary_ curve of 31 nM, which is only a factor of 2.1 off of the measured value. The calculated ternary *K*_d_ *K*_C,2_=*K*_C,1_/α=109 nM/1509=0.7 nM shows that the EC_50_ value of the %C_bound, ternary_ curve is different from *K*_C,2_, i.e., monitoring the binding events through the stronger binding protein does not enable the direct determination of *K*_C,2_ and estimation of the intrinsic cooperativity α would require a mathematical model (Fig. 12 left).

#### R,S-SLF-12: negative control with absolute %A_max_=0%

**R,S-SLF-12** was selected as a negative control compound, which has a high affinity for FKBP12 but does not form any ternary complex according to the TR-FRET assay.

##### Binding study through MAPRE1

The EC_50_ value of the %T_bound, binary_ curve of **R,S-SLF-12** showed no binding to MAPRE1 up to [L_tot_]=200 μM. This negative result indirectly confirms that the above affinities to MAPRE1 are real and required to characterize the cooperativity of the studied ternary complex system. It also emphasizes the importance to use methods capable of accurately detecting very weak affinities (Fig. 13 right).

**Fig. 13.**
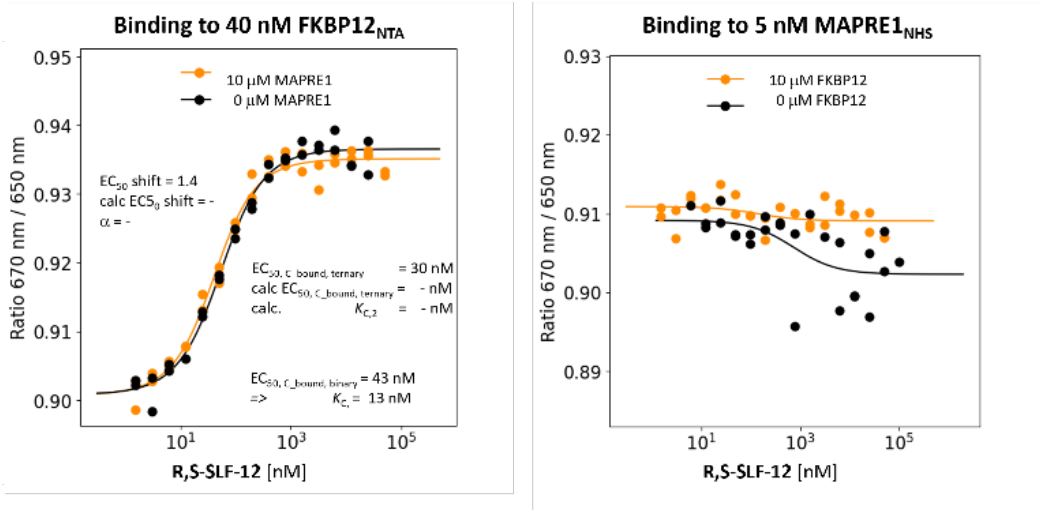
Binding and calculated data for the negative control compound R,S-SLF-12. Right: Neither binary nor ternary binding to MAPRE1 was observed for R,S-SLF-12, which is consistent with the results of the TR-FRET proximity assay (%A_max_=0%). This result indirectly shows that the observed bindings to MAPRE1 are real for the other compounds. Left: To confirm that there is no ternary binding for MAPRE1, no significant shift was observed for the binding of FKBP12. This result confirms the clean correlation between the two directions of binding measurement through MAPRE1 and FKBP12 respectively.

##### Binding study through FKBP12

For the negative control **R,S-SLF-12**, the EC_50_ value of the %C_bound, binary_ curve of 43 nM (i.e., *K*_C,1_=13 nM) to a measured EC_50_ of the %C_bound, ternary_ curve of 30 nM, which results in an apparent cooperativity of 1.4. Since [C_tot_]=40 nM is even higher than the *K*_C,1_=13 nM any *K*_C,2_ lower than *K*_C,1_ (i.e., for α>1) would not be measurable with the given assay setup. Without measurable binding to MAPRE1 and given the lack of a %A_max_ value above background (%A_max_=0%), the observed shift of %C_bound, binary_ of 43 nM to %C_bound, ternary_ of 30 nM by factor 1.4 is too small to be interpreted as a proof for an underlying intrinsic cooperativity α (Fig. 13 left).

#### S,S-SLF-1d: epimeric variant of R,S-SLF-1a with %A_max_ = 0%

**S,S-SLF-1d** is the epimer of **R,S-SLF-1a** and differs only in the stereochemistry at the α-position of the 4-methylenepiperidine-2-carboxyamide unit, which is S instead of R. In the TR-FRET proximity assay, **S,S-SLF-1d** showed no evidence of ternary complex formation (%A_max_=0%) indicating the high specificity of the interactions within the ternary complex FKBP12:**R,S-SLF-1a**:MAPRE1.

##### Binding study through MAPRE1

The EC_50_ value of the %T_bound, binary_ curve of **S,S-SLF-1d** is interpolated to 1.4 mM compared to 2.0 mM for **R,S-SLF-1a**. The same binding experiment but in the presence of 10 μM FKBP12, i.e., [C_tot_]=1428·*K*_C,1_, shifted the EC_50_ value of the %T_bound, ternary_ curve to a [L_tot_] higher than 1.4 mM but it was not possible to identify any *K*_d_. No positive cooperativity (α>1) is observed but negative cooperativity (α<1) cannot be excluded. This would mean that despite the presence of a high excess of FKBP12, **S,S-SLF-1d** is at best bound to MAPRE1 with the same affinity as in the absence of FKBP12. This is a good example of how the strong affinity to the chaperone protein C cannot compensate for the very weak affinity of the target protein T if there is no positive cooperativity α (Fig. 14 right).

**Fig. 14.**
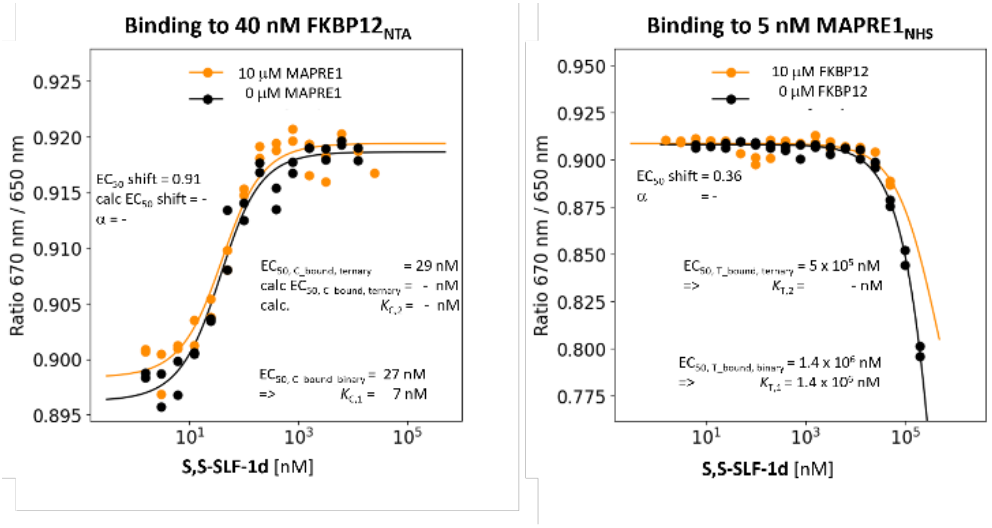
Binding and calculated data for **S,S-SLF-1d**. Right: No binding to MAPRE1 was observed for the negative control compound S,S-SLF-1d, which is consistent with the results of the TR-FRET proximity assay (%A_max_=0%). This result indirectly shows that the observed binding to MAPRE1 is real for the other compounds tested. Left: As a consequence, no significant shift was observed. This result shows the clean correlation between the two directions of the binding measurements for MAPRE1 and FKBP12, respectively.

##### Binding study through FKPB12

EC_50_ of the **S,S-SLF-1d**’s %C_bound, binary_ curve of 27 nM, i.e., *K*_C,1_=7 nM, showed a measured EC_50_ of the %C_bound, ternary_ curve of 29 nM and no apparent cooperativity was observed as the assay was at the bottom with this *K*_C,1_ (Fig. 14 left). This again emphasizes the advantage measuring cooperativity through the weaker binding protein.

#### Measuring native (intrinsic) protein–protein interactions

Intrinsic affinities between the two proteins C and T, i.e., without the presence of a molecular glue add a third pathway that enables the ligand to bind to the chaperone–target complex. In this case, the ternary complex-forming equilibrium would require additional constraints in the mathematical model and impact on the determination of the intrinsic cooperativity α. Importantly, in our study no intrinsic interactions between MAPRE1 and FKPB12 are measurable (Fig. 15) under the current experimental conditions (40 nM FKBP12 vs. 10,000 nM MARPRE1 or 5 nM MAPRE1 vs. 10,000 nM FKBP12) and confirms the observation from NMR studies.^30^

**Fig. 15.**
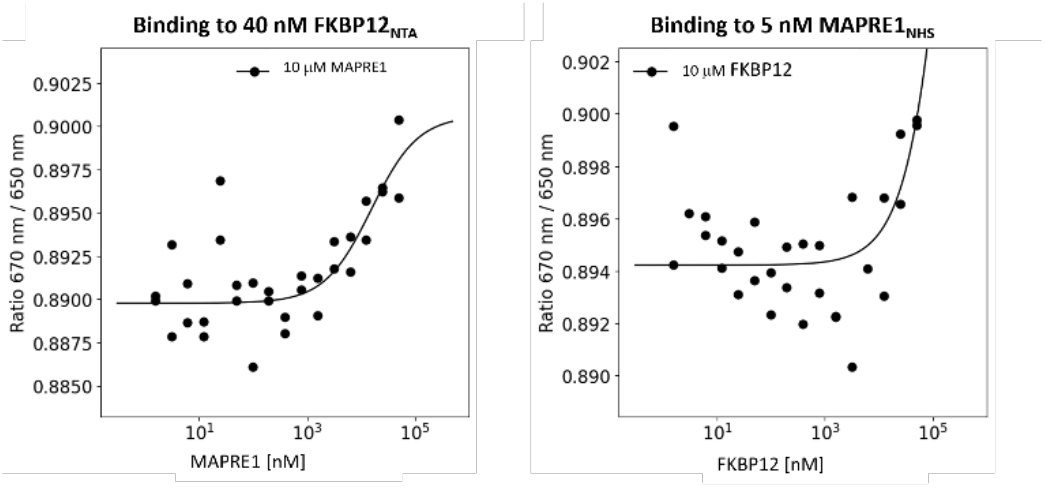
Measuring native protein-protein interaction. The left-hand side fit that can be interpreted as non-binding since the observed amplitude is too small (note the fine scale of the y-axis). This may either reflect gaussian noise or very weak binding with protein affinities much greater than 100 µM. From the right-hand side graph that displays the same interaction but in a reversed fashion no binding is indicated, which suggests no MAPRE1–FKBP12 interaction (see also NMR experiments in our previous work).^**30**^

## Discussion

### General remarks

In our previous publication^29^ we proposed a workflow to retrieve cooperativity α from three measured values for known [C_tot_] and [T_tot_], namely:

a. the EC_50_ value of the %C_bound, binary_ curve to determine *K*_C,1_
b. the EC_50_ value of the %T_bound, binary_ curve to determine *K*_T,1_
c. the EC_50_ value of the %T_bound, ternary_ or %C_bound, ternary_ curve.

We have further described that monitoring by the weaker binding protein produces the larger and thus more accurate apparent cooperativity. Moreover, the greater the excess of the counter protein, the closer the apparent cooperativity and intrinsic cooperativity α. This implies that the EC_50_ value of the %T_bound, ternary_ (or %C_bound, ternary_) curve approaches *K*_T,2_ (or *K*_C,2_) until the minimal offset is reached, which refers to 50% of the concentration of the protein that is used to monitor the binding.

If the numerical difference in affinity to either of the two proteins is not large, i.e., if the ternary complex-forming compound is a bifunctional compound rather than a molecular glue, the previously presented comprehensive mathematical model^29^ describes all simultaneously occurring equilibria under non-saturation conditions and enables to determine the intrinsic cooperativity α. If ternary complexes are formed from molecular glue-like compounds, the difference in affinity between C and T becomes large to very large. If, in addition, a large excess of C can be applied experimentally this results in the following “saturation conditions”:

a. practically no TL complex can be formed,
b. the entire ligand is bound to C into CL,
c. [L_tot_] corresponds to [CL]. For the species that is recruited to T, the EC_50_ value of the %T_bound, ternary_ curve coincides with *K*_T,2_ or at least approaches it (until the offset of half the target protein is reached).

The aim of a binding study with a small number of molecular glues that form a ternary complex with FKBP12 and MAPRE1 was to demonstrate the practical value of the above-mentioned relationships. The selection of compounds for the binding study was based on the %A_max_ values from the TR-FRET proximity assay. %A_max_ values represent a relative measure of the amount of ternary complex formed regardless of whether the driving force for the formation of the ternary complex originates from independent binary affinities to either two proteins or from an underlying cooperativity α. When saturation conditions are achieved, a mathematical model is not required and enables the direct retrieval of the intrinsic cooperativity α by monitoring the binding with only one binding assay. A prerequisite for this is the availability of a method that can measure weak to very weak ligand binding to proteins, preferably as a target engagement assay, i.e., not as a competitive binding assay. For our work, we chose the spectral shift method to measure binding to FKBP12 and MAPRE1 as this method only requires labelling of the monitoring protein and offers high sensitivity.

We are aware that our study consists of only a small number of compounds and that the accuracy of the measured affinities becomes worse with increasing *K*_d_s. However, we believe that the approximation of the weak binary affinities relative to each other should show the same bias. This means that although a more precise measurement would be desirable, it is not experimentally feasible and therefore should not significantly change the determined ranking of cooperativity and its magnitude.

### Correlation of %A_max_ with intrinsic cooperativity α

Changes in the intrinsic cooperativity α alone do not affect the required ligand concentration for maximizing the formation of ternary complexes if *K*_C,1_ and *K*_T,2_ remain the same, but they do affect the amplitude of the curve, i.e., the higher α is, the higher the concentration of the ternary complex ([CLT]) at a given [L_tot_]. In contrast, changes in *K*_C,1_ and *K*_T,2_ at a constant intrinsic cooperativity α shift the required ligand concentration [L_tot_] for the maximum of the curve *and* its amplitude. Changes in the %A_max_ values of a proximity assay together with a shift in the maximum of the CLT formation curve may therefore be the result of a combination of changes in cooperativity α and changes in *K*_C,1_ and/or *K*_T,1_ or just the latter alone. That is, in general, it is not possible to attribute changes in %A_max_ values (in combination with shifts) to changes in any binary affinity and/or intrinsic cooperativity α without a mathematical model that can handle unsaturated conditions. Only if it were possible to keep *K*_C,1_ and *K*_T,2_ constant and only change the intrinsic cooperativity α, changes in the %A_max_ value could be unambiguously assigned with changes of α. However, this is rarely the case. A compound with a different intrinsic cooperativity α generally also shows different binary affinities for the two proteins. If the correctly and independently measured intrinsic cooperativity α and the %A_max_ values for the analogs indeed show a trend correlation, as is the case for the series in this work, this is retrospectively strong evidence that ternary complex formation is driven by the increase in cooperativity α rather than the increase in binary affinities *K*_C,1_ and/or *K*_T,1_. In the described series of molecular glues with measurable %A_max_ values in the TR-FRET proximity assay, the values for *K*_C,1_ range from 21–234 nM, which corresponds to about a factor of 10. The compound with the weakest binary affinity has the highest intrinsic cooperativity α and the highest %A_max_ value, suggesting that there is indeed a correlation between the two values. The values of *K*_T,1_ vary from 0.5– 2.0 mM (4-fold). The values of cooperativity α range from 237 to 10,000 (factor 42) resulting in a trend correlation and underlines for this series that the increase in thermodynamic stability is due to an improvement in cooperativity α (Table 1, column 3 & 4, Fig. 16).

**Fig. 16.**
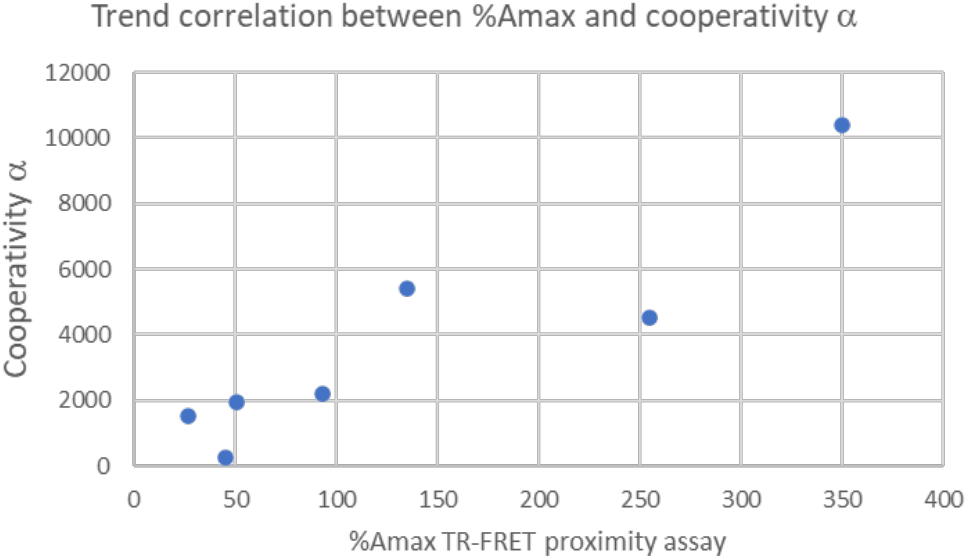
The trend correlation between cooperativity α and the previously reported %A_max_ values confirm that for the studied series of compounds, an increase in the concentration of ternary complexes (indicated by the increase in %A_max_ values) is driven by an increase in cooperativity α.

### Correlation between [C_tot_]/*K*_C,1_ and the ratio of %T_bound, ternary_ over *K*_T,2_

Above, we discussed the influence of the excess of the chaperone protein on whether the EC_50_ of the %T_bound, ternary_ curve coincides with *K*_T,2_ or not. It became clear that “[C_tot_] greater than 100·*K*_C,1_” is a good dump rule to ensure that all ligand L that is not bound into CLT is bound into CL. The more this is the case, the more the value of the EC_50_ of the %T_bound, ternary_ curve subtracted by [T_tot_]/2 approaches *K*_T,2_ because then [L_tot_] corresponds directly to the “ligand” concentration CL. Since in a series of binding experiments [C_tot_] is normally kept constant over a range of compounds tested, but *K*_C,1_ varies from compound to compound, the ratio of [C_tot_]/*K*_C,1_ also varies (Table 1 column 7). In the series described ratios between 41 (Table 1, column 7, entry 1) and 476 (entry 5) occur, which directly determines the degree of saturation, i.e., the degree of coincidence of [L_tot_] with [CL], which allows *K*_T,2_ to be read off from [L_tot_]. In other words, the differences in saturation affect the extent to which all of the ligand that is not bound into CLT is bound to CL, which is directly reflected in the difference between the EC_50_ of the %T_bound, ternary_ curve (minus [T_tot_]/2 for the CLT formed at the inflection point) and *K*_T,2_. Complete saturation leads to no difference, because then [L_tot_] in the EC_50_ corresponds to the concentration of the free ligand in a binary experiment. Fig. 17 shows the correlation of the difference between the measured EC_50_ of the %T_bound, ternary_ curve subtracted by [T_tot_]/2 and the model-based *K*_T,2_ with the ratio of [C_tot_]/*K*_C,1_. Since values on the y-axis (applied protein concentration, measured *K*_C,1_) are independent of the parameters on the X-axis (measured EC_50_ values over calculated *K*_T,2_), the generally good correlation shows the good agreement between the model and the corresponding binding experiments.

**Fig. 17.**
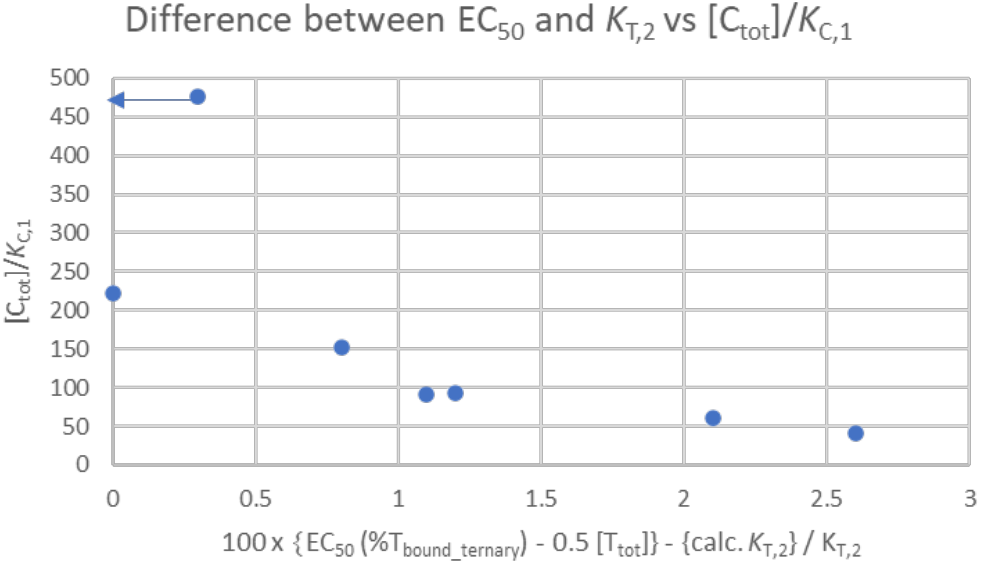
The ratio of [C_tot_]/*K*_C,1_ determines the degree of saturation, which means that at EC_50_ all ligand (=[L_tot_]) that is not bound into CLT (i.e., 50% of [T_tot_]) is bound into CL. Under these conditions, the (normalized) difference between EC_50_–[T_tot_]/2 and *K*_T,2_ approach 0, which is the case for [C_tot_]/*K*_T,2_ values greater than 200. The upper data point displayed at [C_tot_]/*K*_T,2_≈460 should read 0 as x-value (experimental error). The trend correlation shows the good accordance between the model and the experimental data.

### Comparison of binding data measured through MAPRE1 vs. FKBP12

Measurement of apparent cooperativity by the weaker binding protein (called T in this work) causes the larger EC_50_ shifts from the EC_50_ values of the %T_bound, binary_ curve to the %T_bound, ternary_ curve than the measurement by the stronger binding protein (called C in this work) and the corresponding shifts from the %C_bound, binary_ curve to the %C_bound, ternary_ curve. However, the gain in window width for apparent cooperativity competes with the loss in accuracy when measuring weak affinities. Another aspect to consider is the position of the tag. If it is too close to the binding site, it can interfere with the formation of the ternary complex, resulting in a lower apparent cooperativity than would theoretically be expected. If the tag is too far from the binding site, sensitivity may suffer, resulting in a lower signal-to-noise ratio and therefore less accurate EC_50_ shift values. In addition, the interference caused by the tag with the correct folding of the corresponding protein may result in lower apparent cooperativities than would be expected based on obtained data through the counter protein. If possible, it is best to measure cooperativity in both directions, as the two data confirm each other independently.

For the series of compounds and binding experiments described in this work, it became clear that the measurement of binding by MAPRE1 was more optimal than by FKBP12. This can be concluded from the better signal-to-noise ratio of the data measured by MAPRE1 compared to FKBP12, but also from the too small EC_50_ shifts of the %C_bound, ternary_ compared to those of the %T_bound, ternary_ curves. Taking the data measured with MAPRE1 as a reference, the shifts measured with the stronger binding FKBP12 are too small by a factor of about 2 (Table 2, column 6). By modelling the potential conformation of the His-tag with Alphafold^33^, we suspected that the NTD-labelled His-tag might be the cause of the impeded binding events, which is well acknowledge in several enzyme-substrate binding assays^34^ because of its close proximity to the negatively charged MAPRE1 surface (Fig. 18). In contrast, since the N-terminus of MAPRE1 is far away from FKBP12, the N-terminal His-tag on MAPRE1 is expected to have a negligible influence in the binding assay.^30^

**Table 2.**
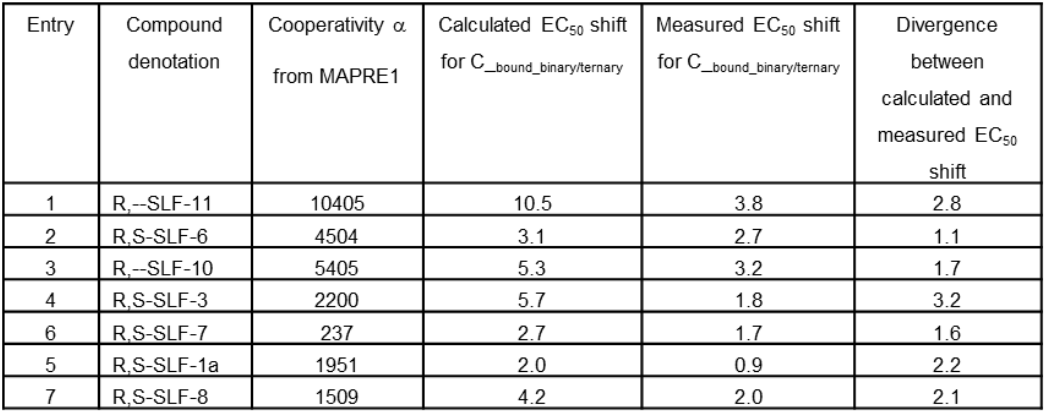
Using the intrinsic cooperativity α values from binary vs. ternary MAPRE1 binding (%T_bound,binary/ternary_) reveals that the calculated EC_50_ shifts for monitoring binary vs. ternary binding through the stronger binding FKBP12 (%C_bound,binary/ternary_) underpredicts these shifts by factor 1.1 to 3.2. Potential reasons include: a) the choice of [C_tot_]=40 nM, so the theoretical bottom of the assay of 20 nM is too close to the calculated EC_50_ values of the %C_bound, ternary_ curve, (b) the location of the tag used for monitoring binding through FKBP12 is compromising ternary complex formation and results in a lower cooperativity α.

**Fig. 18.**
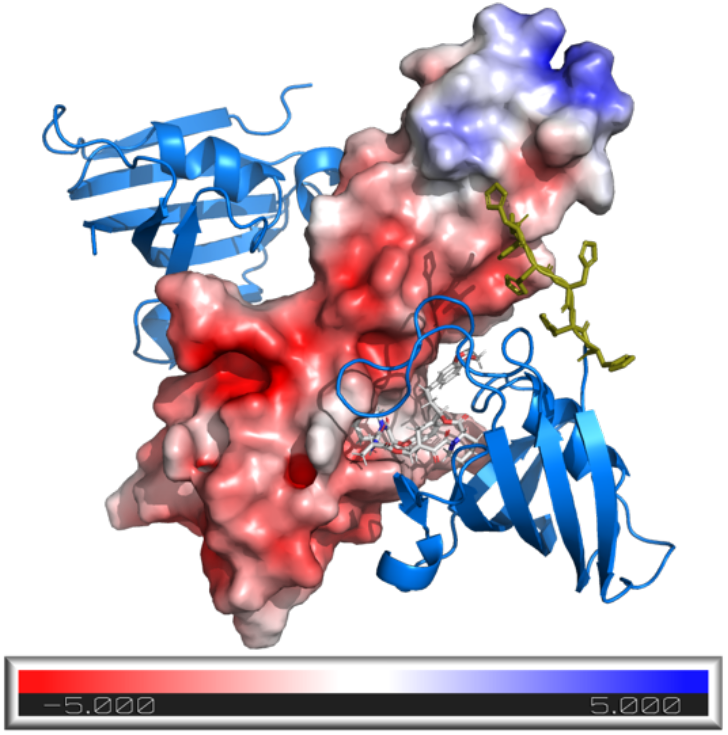
Potential interactions between the His-tag (dark yellow licorice) of FKBP12 (blue cartoon) and MAPRE1 (surface) in presence of the molecular glue R,S-SLF-8 (white licorice). The electrostatic potential of MAPRE1 is color-coded in the unit of k_b_T/e_c_ (k_b_ the Boltzmann constant, T the temperature, e_c_ the electron charge).

## Conclusions

Ternary complexes induced by molecular glues retrieve a significant proportion of their thermodynamic stability through interactions that occur exclusively within the formed ternary complex. These interactions lead to an enhanced ligand binding affinity of the ligand compared to the affinity of the molecular glue to either of the two proteins alone. In general, it is not possible to quantify the degree of this contribution, termed intrinsic cooperativity α, without a comprehensive mathematical model that accounts for all events under non-saturating conditions. This publication describes a series of conditions that allows the determination of the intrinsic cooperativity α from a single binding assay alone, without the need for mathematical modelling. These conditions are fulfilled when the apparent and the intrinsic cooperativity α coincide. We also formalized the relationship between the values of *K*_d_ and EC_50_ as a function of the applied protein concentrations. We have emphasized the value of experimental methods to determine weak binding affinities, but also pointed out ways out when such methods are not available. In series of forthcoming publications, we will present the results of molecular dynamics (MD) studies that rationalize the cooperativities described in this publication from a structural point of view.

## Data availability

The data supporting this article have been included as part of the Supplementary Information of this and of our previous publication.^30^

## Acknowledgements

We thank Pablo Gabriel for help with formatting and Greg Hollingworth for careful proofreading and improvement of the manuscript.

