## Supplemental information for "Experimental conditions to retrieve intrinsic cooperativity α directly from single binding assay data exemplified by the ternary complex formation of FKBP12, MAPRE1 and macrocyclic molecular glues"

**Parameters under which cooperativity α can be determined without a mathematical model and only by one binding assay, using the example of ternary complexes between FKBP12, MAPRE1 and macrocyclic molecular glues**

Jan Schnatwinkel[1], Richard R. Stein[2], Michael Salcius[4], Julian Wong[5], Shu-Yu Chen[6], Marianne Fouché[3], Hans-Joerg Roth*[3]

[1]Nanotemper Technologies GmbH, Tölzer Str. 1, 81369 Munich, Germany

[2]Novartis Pharma AG, CH-4056 Basel, Switzerland

[3]Biomedical Research, Novartis Pharma AG, Fabrikstrasse 2, CH-4056 Basel, Switzerland

[4]Novartis Biomedical Research, 181 Massachusetts Ave, Cambridge, USA

[5]Novartis Biomedical Research,----San Diego, USA

[6]Eidgenösisch Technische Hochschule, Inst. Mol. Phys. Wiss, Zürich, Switzerland

### SUPPLEMENTAL INFORMATION

#### Experimentals Spectral Shift binding assay

His-tagged FKBP12 was labeled using the His-Tag Labeling Kit RED-tris-NTA 2nd Generation (MO-L018; NanoTemper Technologies GmbH).[1] The affinity between NTA dye and the His-tag of FKBP12 was determined as 7 nM. MAPRE1 was labeled using the Protein Labeling Kit RED-NHS 2nd Generation (MO-L011; NanoTemper Technologies GmbH).[1] In brief, 10 µM of MAPRE1 were incubated with a 3-fold molar excess of fluorophore for 30 minutes. The unreacted fluorophore was removed by performing a size exclusion purification (B-column). Collecting the protein fraction yielded 1.8 µM labeled MAPRE1, with a degree-of-labeling of 0.8, as determined on a Nanodrop.

A 1:2 dilution series of all compounds (starting from the 10 mM stock concentration) was prepared in pure DMSO. Each concentration was diluted 25-fold with 20 mM HEPES, pH 7.4, 150 mM NaCl, 1 mM DTT, 0.01% Pluronic F-127. Interaction measurements were prepared in duplicates. For this first 10 µL of the respective ligand dilution was transferred into a Dianthus Microwell Plate (DI-P001A). Another 10 µL of 2-fold concentrated labelled protein (either in the absence, or the presence of 20 µM counter protein, both in 20 mM HEPES, pH 7.4, 150 mM NaCl, 1 mM DTT, 0.01% Pluronic F-127) then yielded the final assay concentrations. For binary interactions of compounds with MAPRE1 a final target concentration of 5 nM and a highest ligand concentration of 50 µM was prepared. For ternary interactions of compounds and MAPRE1 in the presence of 10 µM unlabelled FKBP12 the highest ligand concentration was increased to 200 µM, the highest concentration achievable while maintaining a constant 2% DMSO concentration. For all interactions measured via FKBP12 the target was prepared at a final concentration of 40 nM, supplemented with 5 nM NTA fluorophore. Both binary and ternary interactions (supplemented with 10 µM of unlabelled MAPRE1, again both in 20 mM HEPES, pH 7.4, 150 mM NaCl, 1 mM DTT, 0.01% Pluronic F-127) were prepared with a highest ligand concentration of 50 µM. After 30 minutes incubation at room temperature in the dark, all interactions were studied on a Dianthus instrument from NanoTemper Technologies GmbH (Munich, Germany) at 25°C. The resulting ratio (F670 nm / F650 nm) was plotted against compound concentration and analyzed in the DI.Screening Analysis software.

#### Molecular Modeling

The His-tagged structure of **FKBP12** is first generated by Alphafold 3[2]and aligned to the FKBP12 protein of the **MAPRE1:R,S-SLF-1a:FKBP12** structure deposited on RCSB (pdbid:XXXX[ref])[1] using pymol 2.5.2 root-mean-square-deviation = 0.036nm with 668 atoms). The electrostatics of the MAPRE1 surface is calculated with the APBS Electrostatics plugin at T = 313K [ref].[3]

1. M. Salcius, A. Tutter, M. Fouché, D. King, A. Dhembi, H. Koc, A. Golosov, W. Jahnke, C. Henry, D. Argoti, W. Jia, L. Pedroc, L. Connor, P. Piechon, R. Denay, E. Sager, J. Kuehnoel, M.-A. Lozach, F. Lima, A. Vitrey, S.-Y. Chen, G. Michaud and H.-J. Roth, *RSC Chemical Biology*, 2024, **in preparation**.
2. J. Abramson, J. Adler, J. Dunger, R. Evans, T. Green, A. Pritzel, O. Ronneberger, L. Willmore, A. J. Ballard, J. Bambrick, S. W. Bodenstein, D. A. Evans, C. C. Hung, M. O'Neill, D. Reiman, K. Tunyasuvunakool, Z. Wu, A. Zemgulyte, E. Arvaniti, C. Beattie, O. Bertolli, A. Bridgland, A. Cherepanov, M. Congreve, A. I. Cowen-Rivers, A. Cowie, M. Figurnov, F. B. Fuchs, H. Gladman, R. Jain, Y. A. Khan, C. M. R. Low, K. Perlin, A. Potapenko, P. Savy, S. Singh, A. Stecula, A. Thillaisundaram, C. Tong, S. Yakneen, E. D. Zhong, M. Zielinski, A. Zidek, V. Bapst, P. Kohli, M. Jaderberg, D. Hassabis and J. M. Jumper, *Nature*, 2024, **630**, 493-500.
3. E. Jurrus, D. Engel, K. Star, K. Monson, J. Brandi, L. E. Felberg, D. H. Brookes, L. Wilson, J. Chen, K. Liles, M. Chun, P. Li, D. W. Gohara, T. Dolinsky, R. Konecny, D. R. Koes, J. E. Nielsen, T. Head-Gordon, W. Geng, R. Krasny, G. W. Wei, M. J. Holst, J. A. McCammon and N. A. Baker, *Protein Sci*, 2018, **27**, 112-128.
